# Genome-wide sequence information reveals recurrent hybridization among diploid wheat wild relatives

**DOI:** 10.1101/678045

**Authors:** Nadine Bernhardt, Jonathan Brassac, Xue Dong, Eva-Maria Willing, C. Hart Poskar, Benjamin Kilian, Frank R. Blattner

## Abstract

Many conflicting hypotheses regarding the relationships among crops and wild species closely related to wheat (the genera *Aegilops*, *Amblyopyrum*, and *Triticum*) have been postulated. The contribution of hybridization to the evolution of these taxa is intensely discussed. To determine possible causes for this, and provide a phylogeny of the diploid taxa based on genome-wide sequence information, independent data was obtained from genotyping-by-sequencing and a target-enrichment experiment that returned 244 low-copy nuclear loci. The data were analyzed with Bayesian, likelihood and coalescent-based methods. *D* statistics were used to test if incomplete lineage sorting alone or together with hybridization is the source for incongruent gene trees. Here we present the phylogeny of all diploid species of the wheat wild relatives. We hypothesize that most of the wheat-group species were shaped by a primordial homoploid hybrid speciation event involving the ancestral *Triticum* and *Am. muticum* lineages to form all other species but *Ae. speltoides*. This hybridization event was followed by multiple introgressions affecting all taxa but *Triticum*. Mostly progenitors of the extant species were involved in these processes, while recent interspecific gene flow seems insignificant. The composite nature of many genomes of wheat group taxa results in complicated patterns of diploid contributions when these lineages are involved in polyploid formation, which is, for example, the case in the tetra-and hexaploid wheats. Our analysis provides phylogenetic relationships and a testable hypothesis for the genome compositions in the basic evolutionary units within the wheat group of Triticeae.

## Introduction

Different molecular marker types resulted in widely incongruent hypotheses of relationships for the species belonging to the wheat wild relatives (WWR) of the grass tribe Triticeae (Mason-Gamer and Kellogg 1996; Escobar et al. 2011; Bernhardt 2015; Glémin et al. 2019), i.e. the genera *Aegilops*, *Amblyopyrum*, and *Triticum* (van Slageren 1994; Kilian et al. 2011). Thus, despite their economic importance both as crops and as wild species contributing to the continued improvement of wheat, no comprehensive and generally agreed phylogeny for these species is currently available. This hampers the understanding of the evolution of morphological, physiological, and genetic traits, the biogeography of the species and their environmental adaptation, polyploid formation, speciation, and ultimately the search for useful alleles for plant breeding.

Hybridization is an important evolutionary process (Mallet et al. 2016). It describes the crossing of individuals belonging to different species. On the homoploid level, i.e. if no whole-genome duplication is involved, hybridization results in first generation (F_1_) offspring that possesses half of the genome of each of its parents. If this F_1_ generation becomes reproductively isolated from its parents and evolves into a new species the process is termed homoploid hybrid speciation. If over time repeated backcrossing with one parent dilutes the contribution of the second parent this process is called introgression and means that genomic material (nuclear, chloroplast or mitochondrial DNA) can cross species borders. In contrast, incomplete lineage sorting (ILS) describes the process where during speciation DNA polymorphisms occurring in an ancestral taxon, are stochastically passed on to daughter taxa. Depending on the allele composition in individuals at certain genomic loci, phylogenetic analyses can arrive at different species relationships when different individuals and/or loci are analyzed (Maddison 1997). As ILS mostly depends on population sizes together with mutation rates, the process of lineage sorting can be modeled in a coalescent framework (Kingman 1982). Although it is not always possible to discern hybridization from ILS, multi-locus coalescent analyses including multiple individuals per species can in part overcome this problem (Green et al. 2010; Durand et al. 2011; Pease and Hahn 2015; Yu and Nakhleh 2015; Solís-Lemus and Ané 2016; Wen and Nakhleh 2018; Chao Zhang et al. 2018).

The recent advent of genomic data for *T. aestivum* (International Wheat Genome Sequencing Consortium 2014, 2018), an allohexaploid with three subgenomes (termed **A**, **B**, and **D**), and the related diploid species *Ae. tauschii* (Jia et al. 2013; Luo et al. 2013, 2017) and *T. urartu* (Ling et al. 2013), allows for the comparative analyses of genome structure and gene content. Marcussen et al. (2014), when analyzing relationships among the three subgenomes of wheat, postulated that the **D**-genome lineage, occurring in *Ae. tauschii*, is of homoploid hybrid origin involving the ancestors of the **A** (occurring in *T. urartu*) and **B** genomes (similar to *Ae. speltoides*). This finding spurred a discussion regarding a hybrid origin of *Ae. tauschii* (Li et al. 2015a, b; Sandve et al. 2015). El Baidouri et al. (2017) analyzed sequences of homeologous genes and transposable elements derived from *T. aestivum* (**ABD**), tetraploid *T. durum* (**AB**), *T. urartu* (**A**), *Ae. speltoides* (**B**), and *Ae. tauschii* (**D**). They deduced that about six million years ago (Mya) an ancestral **D** genome introgressed into a homoploid hybrid of the ancestral **A** and **B** genomes. The ancestral **D** genome went extinct sometime later. Today’s **D** genome, occurring in diploid *Ae. tauschii* and as one subgenome in *T. aestivum* and other polyploid species of *Aegilops*, is, therefore, a hybrid genome combining three genomes (El Baidouri et al. 2017). As the **B** genome of polyploid wheat is different from its closest extant relative *Ae. speltoides*, they assumed that the **B** genome itself might also have been introgressed by species of the **S** genome group of *Aegilops* sect. *Sitopsis*. Recently, Glémin et al. (2019) developed a new framework to investigate hybridizations. Based on transcriptome data for all species, they proposed a complex scenario of hybridizations identifying *Am. muticum* (**T**), instead of *Ae. speltoides* (**B**), as an ancestor of the **D**-genome lineage and at least two more hybridization events.

In Triticeae it is generally agreed that the diploid taxa and cytotypes form the basic units of evolution and are involved in different combinations in the formation of polyploid taxa (Kellogg 2015). Polyploids occur mostly as allopolyploid taxa combining the genomes of different parental species after hybridization and whole-genome duplication (WGD). Except for Glémin et al. (2019), the recent studies of the evolution of wheat included only a few species and mostly single individuals (although with huge amount of genome data) of wheat wild relatives. Here we describe the analyses of two genome-wide datasets obtained for all diploid species of *Aegilops*, *Amblyopyrum*, and *Triticum* and always multiple individuals per taxon to improve the understanding of evolutionary relationships in the wheat group. This work employs DNA sequences of 244 nuclear low-copy genes uniformly distributed among all seven chromosomes of the taxa. These were obtained through a set of gene-specific hybridization probes used to enrich the target loci prior to next-generation sequencing (Hyb-seq; Weitemier et al. 2014). Based on this set of genes, species relationships were calculated using diverse phylogenetic algorithms. In addition, genome-wide single-nucleotide polymorphism (SNP) data was obtained through genotyping-by-sequencing (GBS; Elshire et al. 2011). Both datasets were compared for signals of directed introgression and hybridization. Our results provide species relationships within the wheat group taxa, and lead to new hypotheses on far-reaching hybridization and introgression influencing the evolutionary origins and composition of all extant basic diploid genomes in this species group.

## Results and Discussion

### Sequence assembly of the target-enriched loci

Loci for target-enrichment were selected via the comparison of available genome information from different Poaceae like *Brachypodium distachyon*, rice and sorghum, barley and wheat (Vogel et al. 2010; Matsumoto et al. 2011; Mayer et al. 2011), aiming for orthologous loci with an even distribution on the genome (SI Materials and Methods). Our design of capture probes was finally based on 451 loci evenly distributed over the **A**, **B**, and **D** genomes of *T. aestivum* (Table S1, Figure S1).

Target-enrichment and Illumina sequencing resulted in 140 million raw reads and 116 million reads after quality filtering. On average 6% of the reads mapped to the chloroplast genome. Of the 451 loci, 25 (5%) were not sufficiently captured (i.e. not captured in most taxa) and were excluded from further analyses. The capture efficiency was usually taxon/accession independent, indicating no (strong) influence of probe design on the capture efficiency (Table S1, Table S2). The sequences retrieved for the 426 well captured nuclear loci were combined into multiple sequence alignments. Visual inspection of these alignments often showed genus-or species-specific patterns of ambiguous positions. Allelic diversity is assumed to be much lower than 1%. This threshold was set based on a comparison with Jakob et al. (2014) that reported an allelic diversity clearly lower than 1% for the analysis of six single-copy loci of large populations of *Hordeum vulgare* subsp. *spontaneum*. Thus, single-copy loci of heterozygous individuals can be expected to show noticeably less than 1% of ambiguous positions in assembled sequences. Since sequenced accessions within a species mainly share the same combinations of polymorphic positions, this points to the existence of paralogous gene copies for a locus, either functional or as pseudogenes, rather than to heterozygous loci. The proportion of ambiguous positions per accession and locus was estimated (Table S3). An average of more than 1% of ambiguous sites in more than five species was detected for 62 (~15%) captured loci. These loci were considered as mainly multi-copy and excluded from further analyses. Moreover, very short or not variable loci were excluded. The median of the mean coverage for the 244 remaining loci was 25X. Large deviations in the mean coverage result from the actually achieved sequencing depth (Table S4a). The loci used for phylogenetic inference had on average a length of 2,278 bp, 43% of non-variable sites and a pairwise-identity of 88% (Table S4b). Concatenation of the 244 nuclear loci in a supermatrix resulted in an alignment with a total length of 555,543 bp.

### Phylogenies based on target-enrichment data

#### Supermatrix approach

The first step of our analysis procedure was to use DNA sequences of nuclear genes enriched through hybridization probes for Illumina sequencing to infer phylogenetic relationships from quality filtered alignments. In addition to the wheat group taxa, we included four diploid species as outgroups representing the barley genus *Hordeum* (Table S5). Maximum likelihood (ML) and Bayesian phylogenetic inference (BI) of the concatenated DNA sequences of all loci (i.e. creating a supermatrix with 555,543 alignment positions) resulted in the phylogenetic relationships provided in Figure S2. In this tree *Ae. speltoides* and *Am. muticum* form a clade that is sister to all other taxa analyzed. Within the latter, *Triticum* is sister group of the remainder of *Aegilops* species. When analyzing the same dataset with maximum parsimony (MP), *Triticum* and *Ae. speltoides*/*Am. muticum* exchange their respective positions in the phylogenetic tree (Fig. S3).

#### Coalescent-based phylogenetic inference

As data concatenation could potentially result in strong support for wrong species relationships (Xi et al. 2015), gene trees were used to infer a coalescent-based species tree. Individual ML gene trees were used as input for ASTRAL (Mirarab et al. 2014; Chao Zhang et al. 2018), which models ILS under the multispecies coalescent (MSC) model (Degnan and Rosenberg 2009) to deduce species relationships. The resulting phylogeny places *Triticum* as sister to *Amblyopyrum* and all *Aegilops* species (Fig. 1A and S4), a topology similar to the one found by MP analysis of the supermatrix (Fig. S3). *Aegilops markgrafii*/*Ae. umbellulata* form a clade with *Ae. comosa*/*Ae. uniaristata* (clade **CUMN**), although with very low statistical support (Fig. 1A).

**Figure 1.**
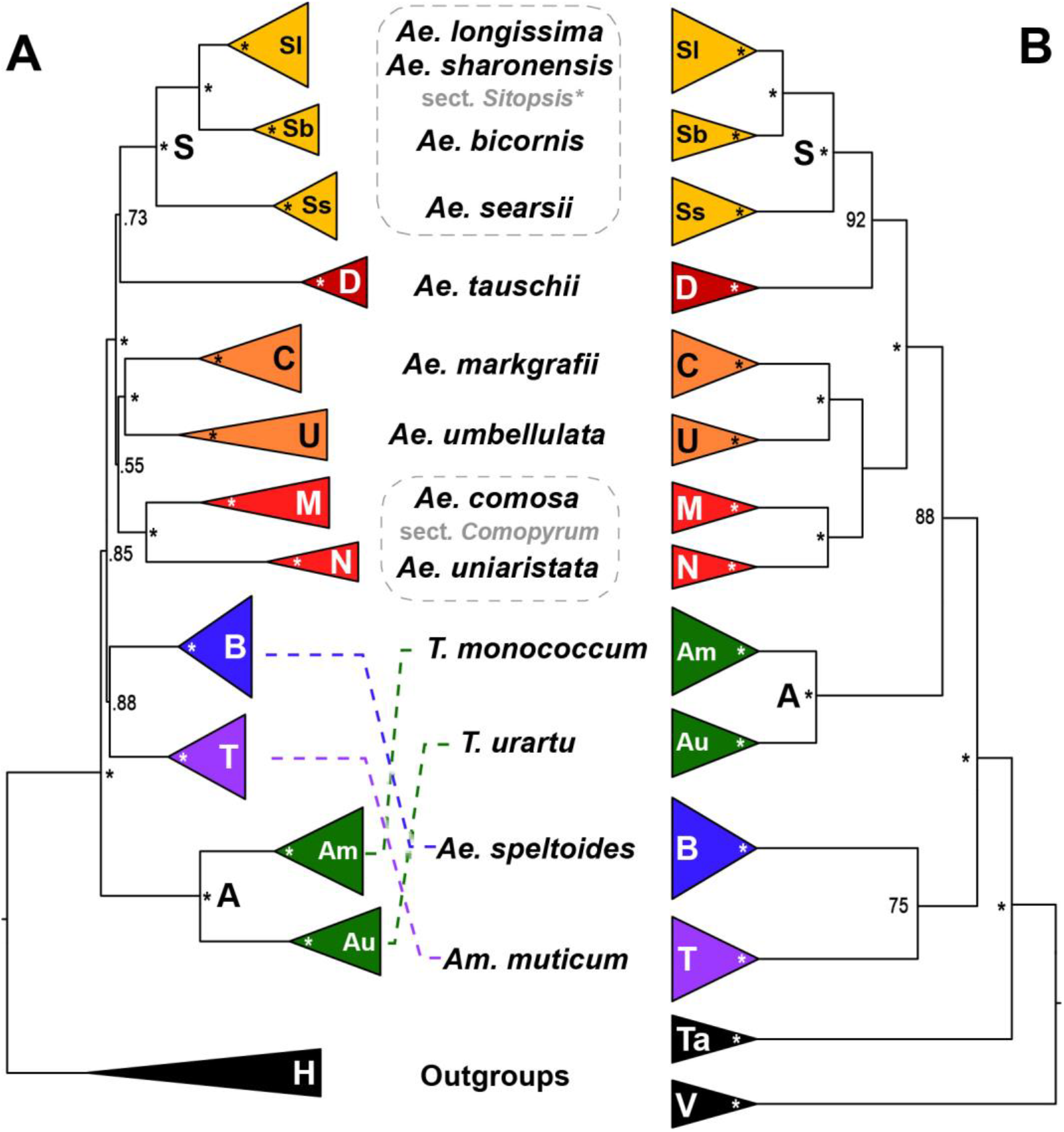
Comparison of coalescent-based phylogenetic trees for the diploid wheat wild relatives. Triticeae-specific genome designations are provided for the respective clades. Fully supported nodes are indicated by asterisks. **A** Schematic representation of the multi-species coalescent tree calculated from separate maximum likelihood gene trees of 244 target-enriched low-copy loci using ASTRAL. Numbers at nodes depict local posterior probabilities. **B** Consensus cladogram derived from a TETRAD analysis of GBS data. Numbers along branches are bootstrap support values (%).

While all 244 individual ML gene trees were in conflict to each other and accessions of the same species may be widely scattered in single topologies (data not shown), all supermatrix phylogenetic approaches (Fig. S2, S3), the ASTRAL analysis (Fig. S4), and the unrooted network obtained via SPLITSTREE (Fig. S5) revealed species to be monophyletic. We, therefore, conclude that ongoing gene flow between species is not significantly impacting the data and extant species can be considered as units.

Low support values in the ASTRAL tree (Fig. 1A and S4) correspond to branches with topological differences when comparing to the supermatrix phylogenies indicating conflicting phylogenetic signal. The degree of gene tree/species tree conflict was investigated in detail with PHYPARTS (Smith et al. 2015), as it could also stem from hybridization/introgression instead of ILS. For most clades comprising several species, no major alternative to the ASTRAL topology could be identified (Fig. S6). However, the clades of **CUMN** and **DS** present in the ASTRAL tree were supported by only seven and 20 out of 244 gene trees, respectively. For the former clade, there were five alternative topologies found to be more frequent involving members of the **CUMN** clade together with either *Ae. tauschii* (**D**) or the *Triticum* species (**A**): **UD** with 14 supporting topologies, **CD** 12, **MND** 10, **AU** 9, and **ND** 8. In the case of **DS**, there were 20 alternative topologies that grouped *Ae*. *speltoides* (**B**) instead of *Ae*. *tauschii* (**D**) together with sect. *Sitopsis* (**S**).

In multi-locus analyses, *Ae. speltoides* always forms a moderately supported clade with *Am. muticum* (**T**), and, as in previous studies (e.g. Petersen et al. 2006; Li et al. 2015a; Bernhardt et al. 2017), it is always clearly separated from the other species of *Aegilops* sect. *Sitopsis* (**S**), as well as from the remaining *Aegilops* species. In the following we will use sect. *Sitopsis*\* to indicate that we refer to the **S**-genome group of sect. *Sitopsis* excluding *Ae*. *speltoides* (**B**) that was earlier placed within this group (van Slageren 1994). *Aegilops tauschii* (**D**), although assumed to be either a homoploid hybrid between the **A**- and **B**-genome lineages (Marcussen et al. 2014; Sandve et al. 2015) or the **A**-, **B**-, and **D**-genome ancestors (El Baidouri et al. 2017), results in all our analyses as sister of sect. *Sitopsis\**. This indicates that an **S**-genome progenitor may have played a role in its formation. This close relationship was not previously postulated, although Marcussen et al. (2014) used sequences of the **S**-genome species *Ae. sharonensis* (International Wheat Genome Sequencing Consortium 2014). However, they excluded them from additional analyses, as they assumed *Ae. sharonensis* itself to be a hybrid involving the **B**-genome lineage. Our data show that not only *Ae. sharonensis* is closely related to *Ae. tauschii* but that shared genome parts most probably involve the entire sect. *Sitopsis\**. Although the relationship to the **B** genome was not found in this initial analysis, it clearly indicates a more complex evolutionary history of the *Ae. tauschii* genome and perhaps also that of sect. *Sitopsis\** in comparison to what was heretofore hypothesized.

Although the discordant topologies revealed by PHYPARTS are potentially better resolved by modeling ILS, they may also result from past hybridizations or gene flow among species. Both processes would violate the assumption of the coalescent analysis that only ILS contributes to deviating gene-tree topologies. Therefore, our sequence data were further analyzed to uncover past hybridization and introgression events.

#### Network approach based on gene tree topologies from target-enrichment data

Even though methods to infer phylogenetic networks are under constant development (e.g. (Yu et al. 2011; Yu and Nakhleh 2015; Solís-Lemus and Ané 2016; Wen et al. 2016; Wen and Nakhleh 2018; Chi Zhang et al. 2018), the analysis of multiple loci, individuals, and species while modeling ILS and reticulations remains computationally expensive (Hejase and Liu 2016; Wen et al. 2018). Thus, resource demanding methods such as full maximum-likelihood or Bayesian inference (Yu et al. 2014; Wen and Nakhleh 2018) failed to infer networks from our entire sequence data. We, therefore, used different strategies of data partitioning by reducing the number of individuals or loci. However, these approaches gave incoherent results across replicates (not shown).

Nevertheless, we were able to obtain phylogenetic networks from the 244 gene tree topologies under the multispecies network coalescent (MSNC) using maximum pseudo-likelihood as implemented in PHYLONET (Yu and Nakhleh 2015). We allowed for zero to five reticulations (Fig. S7a-f). If no hybridization was assumed, the tree with the best log pseudo-likelihood (−7,617,218) had a topology similar to the one obtained via ASTRAL (Fig. 1A, S4). However, poorly supported clades were dissolved resulting in a grade with *Triticum* as sister to the rest of the species, *Am. muticum* and *Ae. speltoides* not being monophyletic, and *Ae. comosa*/*Ae. uniaristata* and *Ae. markgrafii*/*Ae. umbellulata* not clustering together. PHYLONET also retrieved the ASTRAL topology among the top five trees with a slightly lower log pseudo-likelihood (−7,617,519). The network with four hybridization nodes (Fig. 2, S7e) was selected with the Akaike information criterion as best-fit. In this network, hybridizations are nested within each other. This suggests a sequence of hybridization events, the first one involves the ancestors of *Am. muticum* and the *Triticum* clade each contributing approximately equal proportions (0.54 and 0.46, respectively) to the common ancestor of all other species except *Ae. speltoides*. This confirms the scenario inferred by Glémin et al. (2019) identifying *Am. muticum* instead of *Ae. speltoides* as one of the genome donors (Marcussen et al. 2014). Sect. *Sitopsis*\* appears as sister to both *Ae. tauschii* and *Ae. markgrafii* and to be introgressed by *Ae. speltoides* (0.31). Finally, the *Ae. comosa*/*Ae. uniaristata* clade is sister to *Ae. markgrafii* with an additional introgression of the *Triticum* clade (0.29). However, phylogenetic networks inferred from gene tree topologies under maximum pseudo-likelihood are not necessarily uniquely encoded by their system of rooted triples and this analysis may return an equivalent network to the true network (Yu and Nakhleh 2015). In this case, the authors suggest investigating the obtained network with other methods and/or data. Here we used GBS to generate genome-wide SNP data from all taxa to evaluate this scenario.

**Figure 2.**
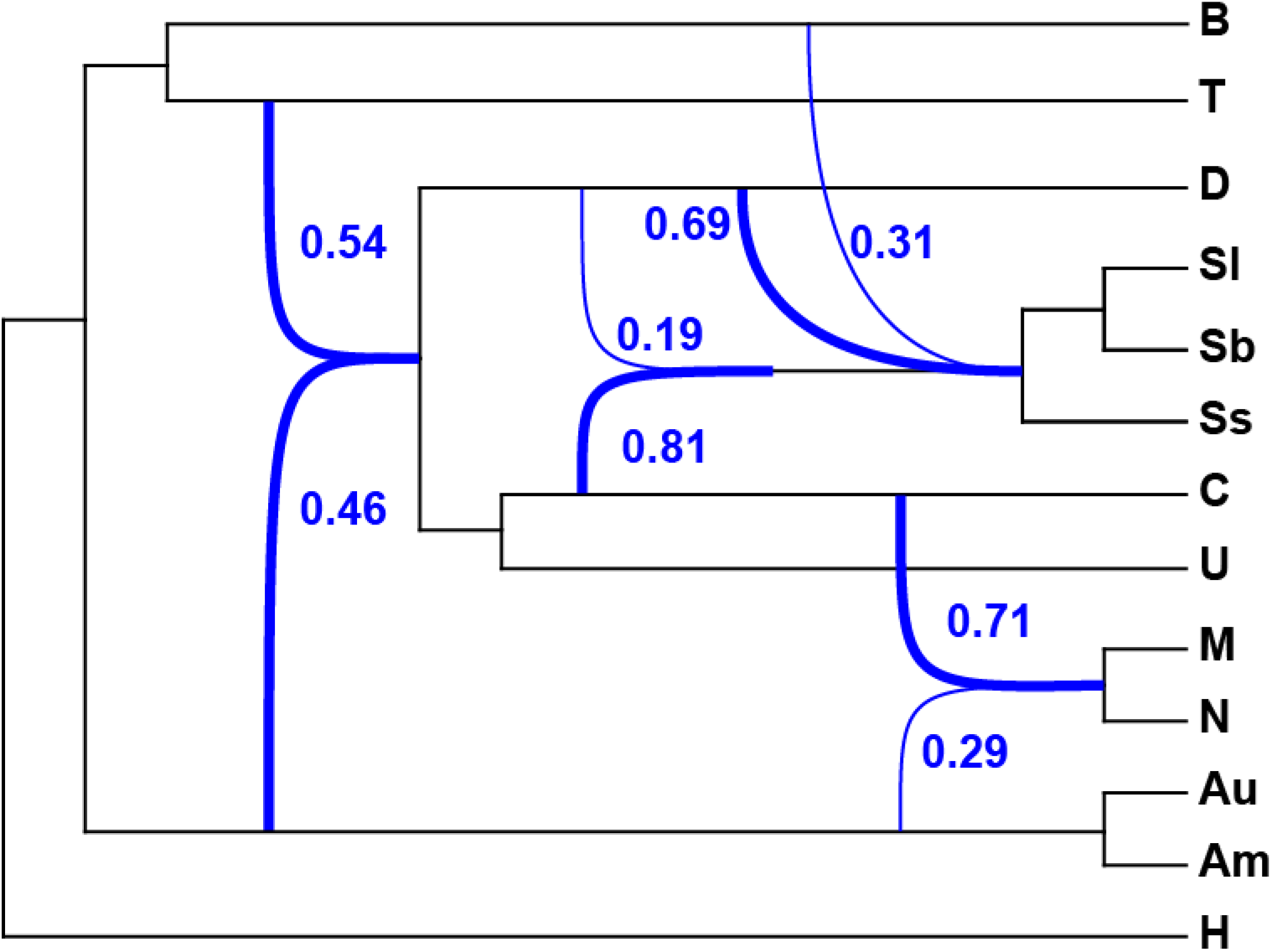
Phylogenetic network inferred under the multispecies network coalescent (MSNC) from the 244 gene tree topologies using maximum pseudo-likelihood. The network with four reticulations was selected as best-fit among zero to five hybridizations calculated with the routine InferNetwork_MPL of PHYLONET under the Akaike information criterion (Fig. S7). Reticulations are indicated by blue arcs with major contribution of species to hybrid lineages indicated by bold lines. Numbers represent estimated inheritance probabilities.

### DNA polymorphisms obtained through genotyping-by-sequencing (GBS)

#### Sequence assembly of the GBS data

To obtain genome-wide SNP data, a two-enzyme GBS analysis (Poland et al. 2012) was performed by cutting the genome with a frequent and a rare-cutting restriction enzyme then sequencing 100 bp of the DNA fragments directly adjacent to the rare restriction sites following Wendler et al. (2014). This method was shown to target the coding parts of the genome (Schreiber et al. 2019). Thus, it can be used to compare SNP patterns between species, which might, in their non-coding genome regions, already be too diverse for meaningful comparisons. As *Hordeum* and the wheat group lineage were already separated 15 Mya (Marcussen et al. 2014), their genomes have diverged substantially. Therefore, we included *Dasypyrum villosum* and *Taeniatherum caput-medusae* as outgroups. These taxa are outside of the wheat group genera (Bernhardt et al. 2017) but still close enough to share multiple GBS loci.

On average 1.65 million reads per sample were obtained from Illumina sequencing. After filtering and clustering on average 222,185 clusters remained per sample. After consensus calling per cluster the number of loci per individual in the assembly was on average 21,000 (with a minimum of 8,472 loci for accession AE 739 of *Ae. speltoides* and maximum of 28,469 loci for accession PI 560122 of *Am. muticum*). In total 140,072 loci having 444,618 phylogenetic informative sites were kept for downstream analysis when specified that at least four individuals had to share a locus (Table S6).

#### GBS-based phylogenetic relationships

To analyze phylogenetic relationships based on the GBS data we conducted an analysis in TETRAD within the IPYRAD package (Eaton 2014; https://github.com/dereneaton/ipyrad). TETRAD uses a single SNP per GBS locus and conducts quartet analyses to infer a species tree that is consistent under the multispecies coalescent. The phylogenetic tree (Fig. 1B, S8) supports the topology of the supermatrix tree of the target-enrichment data (Fig. S2) with respect to the relative positions of *Triticum* and *Ae. speltoides*/*Am. muticum* and of the ASTRAL tree regarding the **MN** and **UC** taxa forming together a weakly supported clade (Fig. 1A). The unrooted phylogenetic network computed by SPLITSTREE (Fig. S9) is concordant with the one for target-enrichment data (Fig. S5) showing that species are monophyletic and can be considered as units for the detection of hybridization.

Even though Zhu and Nakhleh (2018) developed a method (i.e. MLE_BiMarkers) able to deal with more than 50 taxa and four hybridizations using bi-allelic markers under the maximum pseudo-likelihood, we could not process our dataset in a reasonable timeframe (i.e. analyses did not finish within 30 days). We assume that the complexity of the relationships, including putative nested hybridization and introgression events (Fig. 2) complicate the inference of a network from the GBS data. Nonetheless, we assessed hybrid relationships with Four- and Five-taxon *D* statistics. Those methods, based on the frequency of shared polymorphisms between taxa, are less computing intensive.

#### GBS-based D statistics for the detection of hybridization and direction of introgression

Under a neutral model of sequence evolution, and if speciation events occur in rapid succession, ILS should result in similar amounts of shared polymorphisms among species derived from a common ancestor. However, if hybridization is involved, the amount of shared alleles shifts towards the species connected through gene flow in comparison to the background signal contributed by ILS. *D* statistics, also known as ABBA–BABA test (Green et al. 2010a; Durand et al. 2011), is able to discern hybridization from ILS by analyzing allele distribution in three taxa in comparison to an outgroup.

All Four-taxon *D* statistic tests were performed species-wise on unlinked SNPs with the routine Dtrios of DSUITE (Malinsky 2019). First, *D. villosum* was set as outgroup to test if *Ta. caput-medusae* was involved in hybridizations with any members of the WWR (Fig. S10). *Taeniatherum caput*-*medusae* then was used as outgroup for all following tests as no hybridization signal was found. A total of 220 tests were performed of which 64 were significant (*p* value < 0.05 after Benjamini-Yekutieli correction) with *D* statistics ranging between 0.10 and 0.33 (Fig. 3, Table S7). All species were involved in potential hybridizations. The strongest signal revealed a relationship between both *Triticum* species and *Ae*. *markgrafii*/*Ae. umbellulata*, and to a lesser extent *Ae. comosa/Ae. uniaristata* and *Ae. tauschii*. A similar, though weaker, pattern was also found for *Am. muticum*. *Aegilops markgrafii* also showed a strong tie with the members of sect. *Sitopsis\** (**S**). This analysis also confirmed the strong and exclusive relationships between *Ae. speltoides* and the latter.

**Figure 3.**
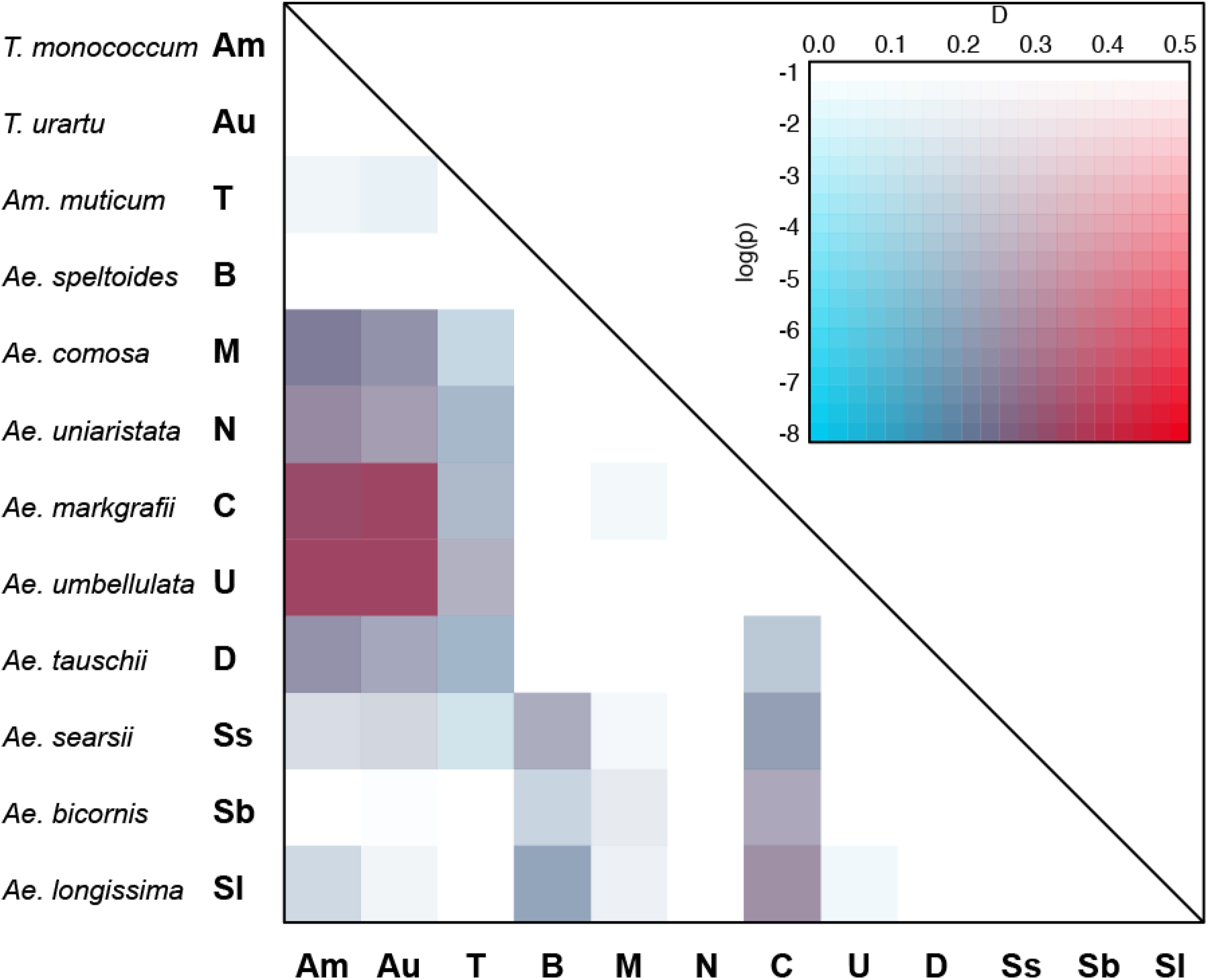
Heatmap summarizing Four-taxon *D* statistic tests using *Taeniatherum caput-medusae* as outgroup. The plot is based on 220 tests. It shows the *D* statistic results and their significance for each pair of species. Red and blue indicate high and low *D* statistic values, respectively. The intensity of the color corresponds to the *p* value (in log-scale) assessed using the block jackknife procedure and corrected with Benjamini-Yekutieli for multiple testing. All *D* statistic results are summarized in Table S7.

An extension of *D* statistics is the *D*_FOIL_ test (Pease and Hahn 2015) that allows not only the detection of hybridization in the presence of ILS but also infers the direction of introgression in a five-taxon phylogeny. This analysis only accepts an alignment of five sequences, therefore we created consensus sequences for each species. *D*_FOIL_ tests were performed with *Ta. caput-medusae* used as the outgroup, to polarize the comparisons of all species. Altogether 216 unique combinations of five taxa were tested but only 143 tests were considered after removing tests that did not fulfill the requirements of estimated divergence times (see Material and Methods; Pease and Hahn 2015). On average 292,602 alignment positions (233,791–379,867) were used resulting in 6,738 (952–10,354) SNP patterns that could be compared (Table S7; Fig. 4). Overall, the relationships inferred are similar to the ones identified by the ABBA–BABA test (Fig. 3; Table S6), however, directions of gene flow could be inferred for nine relationships (11 tests). A large proportion of tests (42) revealed undirected patterns involving three taxa indicative of complex or ancient introgressions, or reciprocal gene flow. Evidence of introgression/hybridization was found for all species (Fig. 4a-k), with a low number of significant tests involving *Ae*. *uniaristata* and *Ae. umbellulata* (Fig. 4e-f) and a high number involving *Ae. markgrafii* and *Ae. longissima* (Fig. 4g and 4k). This analysis confirms the close relationships between the members of sect. *Sitopsis\** (**S**) and *Ae. speltoides* (**B**), but, in contrast to the network inferred with PHYLONET (Fig. 2), *D*_FOIL_ identifies gene flow from **S** to **B** (Fig. 4b). Among the members of sect. *Sitopsis*\*, *Ae. longissima* (**Sl**) appeared as a major introgressor of **B** but also of *Ae. comosa* (**M**), *Ae. markgrafii* (**C**), and *Ae. tauschii* (**D**) (Fig. 4k). This may explain the high number of tests returning undirected signal involving those four species. The close relationship between *Triticum* species and the **CUMND** clade was confirmed although no direction could be inferred (Fig. 4c). This analysis also suggests that *Am. muticum* was affected by gene flow from *Ae. comosa* and *Ae. tauschii* (Fig. 4a).

**Figure 4.**
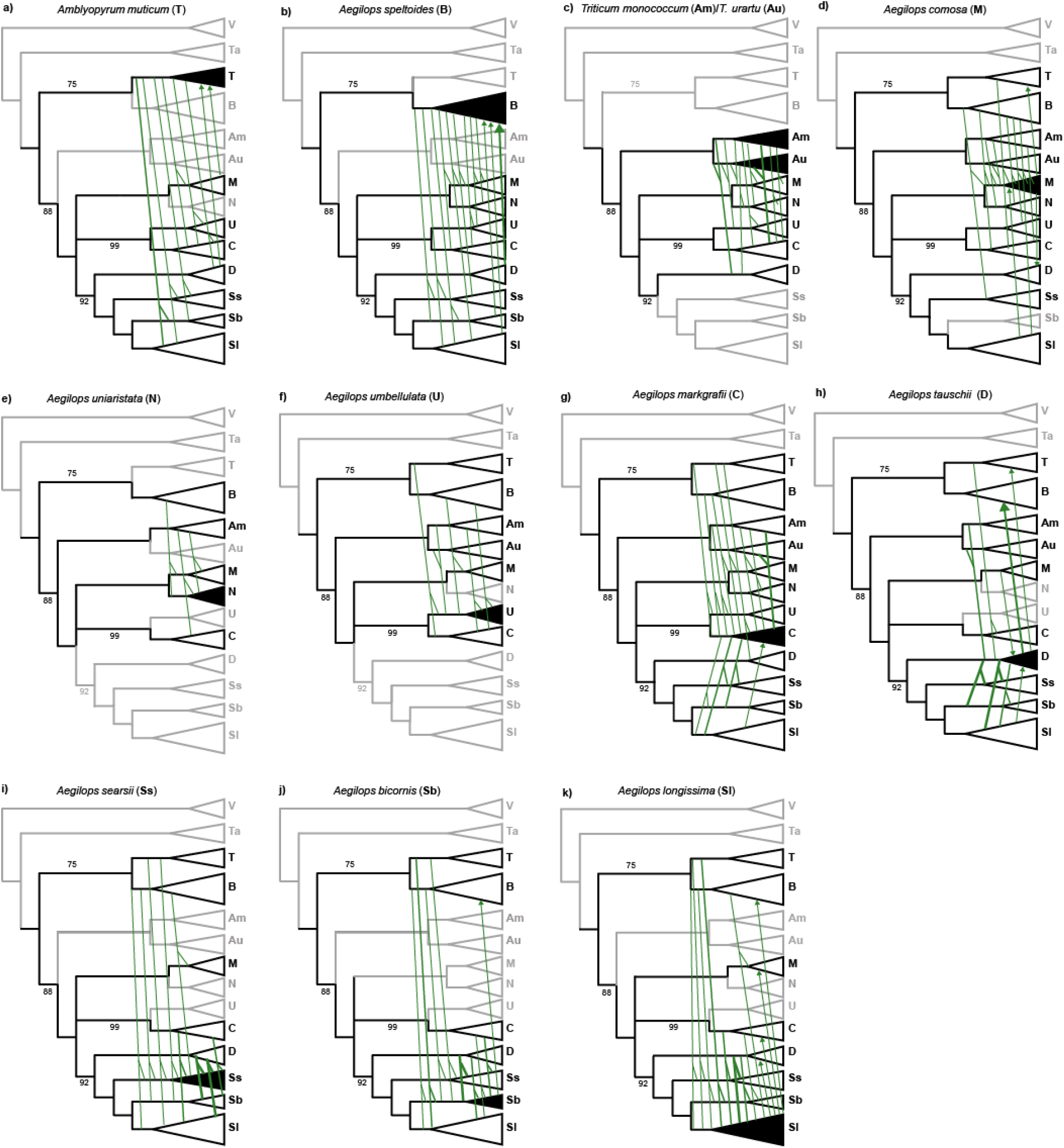
Representation of *D*_FOIL_ results for genotyping-by-sequencing data. All significant relationships after Benjamini-Yekutieli correction are shown on a modified version of the TETRAD species tree. Each tree shows all significant relationships for a focal taxon. An arrow tip indicates the direction of hybridization/introgressions between two taxa. Undirected relationships involving three taxa are shown using a branched line. Taxa not contributing to hybridization signal for the focal taxon are shown in grey for easier visualization. All D_FOIL_ results are summarized in Table S8.

### Homoploid hybrid speciation and major introgressions

In the following, we describe our hypothesis for the evolution of WWR (Fig. 5). Overall, the scenario inferred is similar to the one identified by Glémin et al. (2019). Nonetheless, as we did not focus on identifying the progenitors of the “**D**-genome lineage” we are able to propose a more complete picture. However, as the relationships we identified are highly reticulate, there are partly alternative scenarios possible. We limit our interpretation to the most strongly supported relationships to avoid false positives (Eaton et al. 2015).

**Figure 5.**
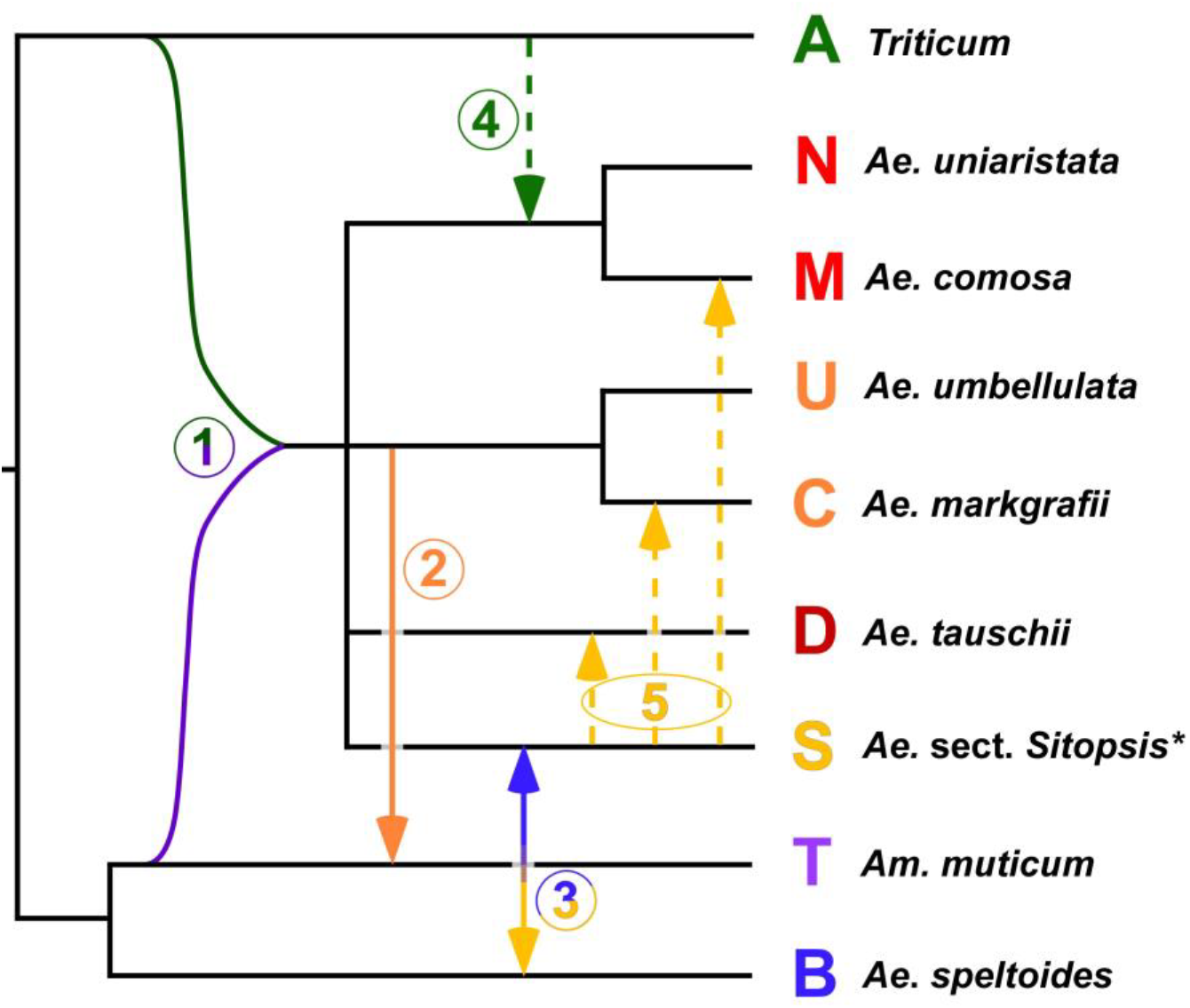
Total evidence evolutionary scenario for the wheat wild relatives. All diploid *Aegilops* species except *Ae. speltoides* are derived from an initial homoploid hybridization event involving the ancient **A** (*Triticum*) and **T** (*Am. muticum*) lineages (1). Strong signals of introgression were found for *Am. muticum* (from the **U**/**C** group; 2) and between *Ae. speltoides* and sect. *Sitopsis* (3). For the latter, introgression seems to have happened in both directions. Weaker signals of introgression (dashed arrows) were found by GBS-based *D* statistics from the *Triticum* (**A**) into the **M**/**N** lineage (4), and (5) from sect. *Sitopsis* into *Ae. tauschii* (**D**), *Ae. markgrafii* (**C**) and *Ae. comosa* (**M**).

As our phylogenetic analyses revealed the monophyly of all species we are certain that hybridizations and introgressions involved mainly ancestral taxa and not the extant species. Our results suggest that there are different groups of taxa, i.e. lineages that introgressed others, lineages that are recipients of introgressions from one or several taxa and/or lineages that originated via homoploid hybrid speciation.

We hypothesize that most of the wheat-group species were shaped by a primordial homoploid hybrid speciation event, i.e. that the *Triticum* lineage merged with the ancestor of *Am. muticum* to form all other species but *Ae. speltoides*. This hybridization event was followed by multiple introgressions affecting all taxa but *Triticum*. In contrast to Glémin et al. (2019), we do not find an introgression of *Triticum* into *Am. muticum*, instead our results indicate that *Am. muticum* may have been introgressed by *Ae*. *umbellulata* or the common ancestor of the **CUMND** clade (Fig. 4a, S7d). Previously published chloroplast phylogenies (Bordbar et al. 2011; Bernhardt et al. 2017) support this event of introgression into **T**, as the chloroplast of *Am. muticum* does not group with *Ae. speltoides* in the chloroplast phylogeny, although both are sister taxa in nuclear phylogenies. These results highlight the pivotal role of *Am. muticum*, instead of *Ae. speltoides* in the formation of the WWR.

For *Ae. speltoides* (**B**) conflicting results were obtained with either sect. *Sitopsis\** (**S**) being introgressed by **B** (Fig. 2) or the other way around (Fig. 4b). This suggests that either reciprocal gene flow occurred between those species or that at least one of the applied methods revealed false positives. Both methods have drawbacks: phylogenetic networks obtained under maximum pseudo-likelihood may not be true but rather equivalent to the true network (Yu and Nakhleh 2015), and *D* statistics are only analyzing three or four taxa simultaneously. Nevertheless, sect. *Sitopsis\**, and especially *Ae. longissima* that has been described as an outcrossing taxon (Escobar et al. 2010), was repeatedly identified as an introgressor as it exhibits relationships with all taxa except the *Triticum* lineage (Fig. 4k).

Signals for the involvement of the sect. *Sitopsis\** genomes can be found in *Ae. comosa* (**M**) and *Ae. markgrafii* (**C**), for which a hybrid origin has been recently proposed (Danilova et al. 2017). Both taxa presented patterns of introgressions different from their respective sister species *Ae. umbellulata* and *Ae. uniaristata*. These two species were involved in the least number of hybridizations. This seems to indicate that **C** and **M** lineages diverged from their sister species due to minor introgressions from *Ae. longissima* or other species of the sect. *Sitopsis\**. It is further suspected that *Ae*. *longissima* or sect. *Sitopsis\** strongly introgressed another, possibly extinct (El Baidouri et al. 2017), member or the progenitor of the **CUMN** clade to form *Ae. tauschii*, as the observed pattern does not resemble a simple sister-species relationship (Fig. 3, 4h, S6). *Aegilops tauschii*, therefore, displays similarities with *Triticum* (**A**), *Ae. comosa* (**M**) and, to a lower extent, *Am. muticum* due to the primordial homoploid hybrid speciation, and is, through its sect. *Sitopsis\** parent, connected to *Ae. speltoides*.

In addition to the major evolutionary scenario developed in this work, past or present gene flow among the different lineages of WWR cannot be ruled out entirely, whenever species come into contact with each other (Arrigo et al. 2011; Bernhardt et al. 2017). The existence of extinct ancestral lineages (Brassac and Blattner 2015) that could not be sampled may, in general, mislead the results of *D* statistics (Beerli 2004; Slatkin 2005). However, in that case *D* statistics are expected to return mostly false-negative test results (Pease and Hahn 2015) instead of arriving at wrong species connections. On the other hand, although we took a conservative approach, ancestral population structure, non-random mating, and small effective population sizes, characteristic of inbreeding species like most wheat wild relative species, could lead to high *D* statistic values (Eriksson and Manica 2012; Martin et al. 2015). New methods accounting for demographic processes at the scale of a genus are necessary to overcome this limitation.

## Conclusions

We obtained DNA sequences of 244 nuclear low-copy genes evenly distributed among the Triticeae chromosomes and genome-wide single-nucleotide polymorphism for all diploid species of the WWR. A combination of different phylogenetic and network approaches together with *D* statistics revealed ancient complex reticulated processes partly involving multiple rounds of introgression as well as at least one homoploid hybrid speciation during the formation of the extant taxa.

Based on our comprehensive taxon sampling we are able to propose a detailed scheme of events that shaped the close relatives of wheat, and is much more complex than previously suggested (Marcussen et al. 2014; Li et al. 2015a, b; Sandve et al. 2015; El Baidouri et al. 2017). With two independent datasets, we were not only able to confirm the scenario developed by Glémin et al. (2019) and that seems to best reflect the evolution of wheat wild relatives but also to uncover more complex pattern of inter-specific gene flow. Our hypothesis is congruent with the proposed formation of the **D**-genome lineage through homoploid hybrid speciation (Marcussen et al. 2014) but proposes, in agreement with Glémin et al. (2019), *Am. muticum* together with the *Triticum* lineage as progenitors. Furthermore, we suggest that *Ae. longissima* or members of sect. *Sitopsis\** played an important role in the formation of *Ae. comosa* (**M**), *Ae. markgrafii* (**C**), and *Ae. tauschii* (**D**). We propose that *Ae. tauschii* belongs to the **CUMN** clade but was introgressed by *Ae. longissima* or sect. *Sitopsis\** thus appearing as its sister species. Moreover, our data provide evidence of gene flow between sect. *Sitopsis\** and the **B**-genome lineage, a hypothesis raised by El Baidouri et al. (2017) and Glémin et al. (2019). We also show that *Am. muticum* cannot be separated from *Aegilops*, as it is sister-taxon to *Ae. speltoides* for nuclear data and is both a progenitor of and introgressed by other *Aegilops* species as shown from *D* statistics and plastid phylogenies (Bordbar et al. 2011; Bernhardt et al. 2017). As the here proposed scenario is highly reticulate, it is necessary to obtain extensive genome information for all diploid species of this group to test predictions regarding composite genomes. Hybrid speciation and introgression should influence genome organization, the presence of syntenic blocks, and the occurrence of different transposable elements within the basic and hybrid lineages of the wheat group taxa. In more general terms the question remains if the important role of hybrid speciation and introgression we found in the wheat group is a peculiarity of these taxa or if it plays an important role in most grasses or generally in plant evolution but was not yet detected, as studies using an approach similar to ours are still mainly in their infancy.

## Materials and Methods

### Plant materials

We analyzed 97 individuals representing all diploid species of the WWR with multiple individuals plus three outgroup taxa (i.e. *Dasypyrum*, *Hordeum*, *Taeniatherum*) of the grass tribe Triticeae (Table S5). All materials were grown from seed and identified based on morphological characters if an inflorescence was produced. Vouchers of the morphologically identified materials were deposited in the herbarium of IPK (GAT). Genome size and ploidy level of 83 individuals were initially verified by flow cytometry and genomic DNA was extracted as in Bernhardt et al. (2017).

### Design of capture probes and library preparation for target-enrichment

We used the assembly of *H. vulgare* cv. ‘Morex’ (Mayer et al. 2012), the only Triticeae draft genome that was available at the time of bait design, to select loci for which orthology could be confirmed when comparing them to the fully sequenced grass genomes of *Brachypodium distachyon*, rice, and sorghum (Vogel et al. 2010; Matsumoto et al. 2011; Mayer et al. 2011). Subsequently, one locus was selected every 0.5 cM on all *H*. *vulgare* chromosomes. These loci were used for BLAST comparisons (Altschul et al. 1990) against available data of *Brachypodium*, rice, sorghum, barley, and wheat. Multiple sequence alignments were built including full-length cDNA (fl-cDNA) and genomic DNA sequences. Finally, 451 loci were chosen for the design of hybridization probes, if they showed (i) a conserved exon-intron structure, (ii) a total length of exonic region larger than 1000 bp with (iii) a minimum size of single exons being 120 bp, and (iv) introns separating adjacent short exons being smaller than 400 bp. The design of capture probes for the selected loci was finally based only on fl-cDNAs from *H. vulgare* and *T. aestivum*, two distantly related Triticeae taxa, and *Brachypodium distachyon*, which was used to broaden the taxonomic spectrum. Capture probes for each of the loci were designed on exon sequences of all three species. The loci used for bait-design are evenly distributed over the **A**, **B**, and **D** genomes of *T. aestivum* (Table S1, Figure S1). The total exonic sequence information considered in bait design amounts to 690 kb. Custom PERL scripts were used to design bait sequences that were submitted to the web-based application eARRAY (Agilent Technologies). A detailed description of the bait design can be found in the Supplementary Information (SI) Material and Methods.

For each of the selected 69 samples (Table S5) 3 µg genomic DNA were sheared into fragments having an average length of 400 bp. The sheared DNA was used in a sequence-capture approach (SureSelect^XT^ Target Enrichment for Illumina Paired-End Sequencing, Agilent Technologies). All samples were barcoded, pooled, and sequenced on the Illumina HiSeq 2000 or MiSeq. For further details see SI Material and Methods.

### Library construction and sequencing for genotyping-by-sequencing (GBS)

GBS and Illumina sequencing were performed for 57 individuals (Table S5) following Wendler et al. (2014). *Dasypyrum villosum* and *Taeniatherum caput*-*medusae* were included as outgroup taxa. For each individual, 200 ng genomic DNA were digested by two restriction enzymes *Pst*I-HF (CTGCAG, NEB Inc.) and *Msp*I (CCGG, NEB Inc.). Sequencing was done on an Illumina HiSeq 2500 obtaining 100 bp single-end reads.

### Target-enrichment data assembly and analyses

#### Assembly

The loci were assembled in a two steps procedure. First, all 451 loci were assembled in a fast and non-stringent approach to evaluate if the capture worked sufficiently and if the loci are truly single-copy in most of the taxa. For each sample, the sequence reads were mapped to the barley genome assembly (Mayer et al. 2012) using the Burrows-Wheeler Alignment (BWA) Tool v. 0.7.8 (Li and Durbin 2009). Consensus sequences were called using SAMTOOLS version 1.1. (Li et al. 2009; Li 2011) and converted into FASTA sequences using VCFUTILS and SEQTK version 1.0 (Heng Li, https://github.com/lh3/seqtk). The percentage of ambiguous sites was determined for each sequence in locus-wise multiple sequence alignments. Allelic diversity is assumed to be much lower than 1% for single- and low-copy-number loci (for comparison see Jakob et al. 2014). Thus, a high percentage of ambiguous positions for sequences of the same species are assumed to reflect the presence of paralogous gene copies. Finally, loci with an average number of ambiguous sites >1% in six or more species of *Aegilops* and *Triticum* were considered as multi-copy (Table S3). Then, the loci found to be mainly low-copy-number loci were kept and selected for a refined assembly procedure if they had a length of at least 1000 bp, contained less than 25% of missing data and at least 15% of parsimony-informative positions, as identified with PAUP*4.0a146 (Swofford 2002). The refined assembly was performed in GENEIOUS v. 10.0.5 (Kearse et al. 2012) as it can reliably assemble short insertions and deletions (Smith 2015). For further details see SI Materials and Methods.

#### Phylogenetic analyses

To infer the phylogeny of the wheat relatives we adopted an analysis approach consisting of the following steps. After aligning the sequences for all loci separately, (i) models of sequence evolution were determined for each locus. (ii) Gene trees were inferred for each locus by maximum likelihood (ML). (iii) The degree of gene tree/species tree conflict was investigated in detail with PHYPARTS. (iv) Concatenated sequences from all loci (supermatrix) were used for Bayesian phylogenetic inference (BI), maximum likelihood (ML), maximum parsimony (MP) and NEIGHBORNET analyses. (v) Multispecies coalescent-based analyses were conducted to infer species trees from the ML gene trees. (vi) Phylogenetic networks were calculated based on the ML gene tree topologies. These analysis steps are detailed below.

#### Gene tree inference

Individual gene trees were inferred using RAXML v. 8.1 (Stamatakis 2014) under the GTRCAT model, rapid bootstrapping of 100 replicates and search for the best-scoring ML tree. To reduce noise from the data, the ML trees were further processed by contracting low support branches (bootstrap values < 10) as suggested by (Chao Zhang et al. 2018) with the Newick utilities function nw_ed and rerooted using the MRCA of *Hordeum* as outgroup with the function nw_reroot (Junier and Zdobnov 2010).

#### Supermatrix phylogeny

Multiple sequence alignments of all 244 loci were concatenated. Bayesian inference was performed in MRBAYES V. 3.2.6 (Ronquist et al. 2012) on CIPRES, Cyberinfrastructure for Phylogenetic Research Science Gateway 3.3 (Miller et al. 2010). The best-fitting models of sequence evolution were estimated by making the MCMC sampling across all substitution models as described in Bernhardt et al. (2017). *Hordeum vulgare* was set as outgroup. An alternative approach to visualize the variation in the data was conducted by computing an unrooted phylogenetic network via SPLITSTREE v. 4.14.8 (Huson and Bryant 2006). The tool was run using the algorithms Uncorrected P, NeighborNet and EqualeAngle for the matrix of the 244 concatenated target-enrichment loci.

An MP analysis of the supermatrix was conducted in PAUP* V. 4.0a146 (Swofford 2002) to see if the phylogeny obtained by BI is sufficiently robust with regards to different analysis algorithms. The MP analysis was run using a heuristic search with 100 random-addition sequences and tree bisection and reconnection (TBR) branch swapping, saving all shortest trees. Node support was evaluated by 500 bootstrap re-samples with the same settings but without random-addition sequences.

#### Coalescent-based species tree estimation

The effect of gene tree conflicts due to ILS was addressed using the short-cut coalescence method ASTRAL (Mirarab et al. 2014; Chao Zhang et al. 2018), which is able to estimate the true species tree with high probability, given a sufficiently large number of correct gene trees under the multispecies coalescent model. ASTRAL-III v. 5.6.3 was run using 244 the ML edited and rerooted gene trees pre-estimated in RAXML.

#### Differences among gene trees

PHYPARTS (Smith et al. 2015) was used to summarize the amount of concordant and conflicting phylogenetic signal from the 244 ML gene trees with the ASTRAL topology as species tree. Visualization of the output was done as in Kates et al. (2018) and Villaverde et al. (2018), and using the phypartspiecharts.py script of M. Johnson available at www.github.com/mossmatters/phyloscripts.

#### Maximum pseudo-likelihood gene tree-based phylogenetic networks estimation

Throughout all analyses *Ae. sharonensis* groups within *Ae. longissima* and *T. monococcum* within *T. boeoticum.* This is in accord with what is already known about these species, i.e. *Ae. sharonensis* and *Ae. longissima* are closely related taxa, and the unified or separate treatment of the two *Triticum* taxa is debated (van Slageren 1994; Bernhardt 2015). Here we use *Ae*. *sharonensis* and *T. boeoticum* if accessions were assigned to this taxon in the donor seed bank. However, due to their strong genetic similarity we treat *Ae*. *sharonensis* and *Ae*. *longissima* as well as *T. boeoticum* and *T. monococcum* con-specific.

The effect of gene tree conflicts due to hybridizations was investigated with the maximum pseudo-likelihood method InferNetwork_MPL (Yu and Nakhleh 2015) included in the package PHYLONET (Than et al. 2008; Wen et al. 2018). The set of ML gene trees analyzed with ASTRAL was used as input for PHYLONET allowing for zero to five hybridizations, other options were left to default. For each analysis, the best network was recorded and they were compared using the Akaike information criterion (AIC; Akaike 1974). As suggested by Yu et al. (2012) and Morales-Briones et al. (2018), the number of parameters was set to the number of branches plus the number of hybridization probabilities being estimated. The network with the lowest AIC score was selected as the best-fit multi-species network. The network was visualized with DENDROSCOPE (Huson and Scornavacca 2012).

### Assembly and analysis of GBS data

The assembly of the GBS data was performed *de novo* using IPYRAD v. 0.7.17 (Eaton 2014; https://github.com/dereneaton/ipyrad), with strict filtering for adapters and restricting the maximum number of heterozygous sites per locus to 25%. Default settings were used for the remaining parameters.

A species tree based on SVDQUARTETS (Chifman and Kubatko 2014) under multispecies coalescence was estimated using TETRAD, as implemented in IPYRAD v. 0.7.17 with 100 bootstrap replicates. For comparison with the target-enrichment data SPLITSTREE v. 4.14.8 (Huson and Bryant 2006) was run using the methods Uncorrected P, NeighborNet and EqualeAngle to compute unrooted phylogenetic networks for 807,909 SNPs of the GBS analysis.

### Identification of hybrid taxa

We used Four-taxon *D* statistics (Green et al. 2010a; Durand et al. 2011; Eaton and Ree 2013) for the GBS data to identify candidate lineages involved in the introgressive hybridization within a fixed phylogeny (((P1, P2) P3), O). Under ILS alone, the number of shared single-nucleotide polymorphisms (SNPs) resulting in an incongruent topology (i.e. ABBA and BABA) are expected to be equivalent. If P3 was involved in an introgressive event with P1, it will share more SNPs with P1 (i.e. BABA patterns), than with P2 (i.e. ABBA patterns).

The VCF file generated by IPYRAD was first filtered with SAMTOOLS/BCFTOOLS (Li 2011) retaining only unlinked SNPs. Four-Taxon *D* statistic tests were performed using the routine Dtrios of DSUITE (Malinsky 2019; https://github.com/millanek/Dsuite). We first tested if *Taeniatherum caput*-*medusae* was involved in any introgressions. As no hybridization signal was found (Fig. S10) and because it is sharing more loci with the WWR than *D*. *villosum*, *Ta. caput*-*medusae* was used as outgroup taxon for all following tests. The VCF file was further processed to exclude all *D. villosum* individuals and DTRIOS was used to perform 220 tests. The ASTRAL topology (Fig. 1A) was used to specify species relationships. *D* statistics significance was assessed using jackknife (Green et al. 2010) on blocks of 100 SNPs. The function *p.adjust* in R 3.5.3 (R Core Team 2019) was used to apply a Benjamini-Yekutieli correction (Benjamini and Yekutieli 2001). All 220 tests are summarized in Table S7. The results were visualized with the Ruby script “plot_d.rb” available from M. Matschiner (https://github.com/mmatschiner).

The *D*_FOIL_ test (Pease and Hahn 2015; https://github.com/jbpease/dfoil/) was used on the GBS data. It relies on a symmetric five-taxon phylogeny (((P1, P2), (P3, P4)), O) to identify the direction of introgressions among the candidate taxa identified using the Four-Taxon *D* statistic. All tests were performed on species-specific consensus sequences. For each species, the alignment of all loci was used to call a consensus sequence that represented all diversity within the species. Therefore, we used the “0% identical” threshold in GENEIOUS that minimizes the number of ambiguities. A custom workflow in GENEIOUS was used to create datasets of five species including *Ta. caput-medusae* as outgroup. For all tests, we made sure that the estimated divergence times fit the assumptions of the program, i.e. that P1 and P2 diverged after P3 and P4 in forward time, by excluding all tests that raised the warning “b” (Table S8). We also used a feature of *D*_FOIL_, i.e. *D*_FOIL_alt, that excludes single derived-allele count for tests with an error warning “c” (Table S8) following Leduc-Robert and Maddison (2018). As a total of 216 tests were conducted, a Benjamini-Yekutieli correction (Benjamini and Yekutieli 2001) was applied to all four statistics for each test with the function *p.adjust* in R 3.5.3 (R Core Team 2019). A significance level of 0.01 was then used on the adjusted *p* values to identify patterns of introgression.

### Contribution of Authors

Designed study: FRB, NB, BK. Coordinated study: NB. Provided data or materials: EMW, BK. Performed experiments: NB. Analyzed data: NB, JB, XD, FRB, and CHP. NB and FRB wrote the initial manuscript. All authors contributed to and approved the final version.

## Supporting information

Table S1 Loci selected for TE

Table S2 TE data read statistics

Table S3 Paralogs identification

Table S4 TE loci statistics and MSAs

Table S5 Plant material

Table S6 GBS data read statistics

Table S7 Four-Taxon D-statistics

Table S8 Five-Taxon D-statistics (Dfoil)

## Acknowledgments

This work was supported by the German Research Foundation (DFG) through grant BL462/10 to FRB and BK, and by basic funds from IPK Gatersleben. We would like to thank M. Pfeiffer for help during the initial steps of bait design and K. Schneeberger for helpful discussions regarding locus choice and data assembly for the target-enrichment experiment. We are grateful to R. Brandt and A. Himmelbach for performing the Illumina sequencing, C. Koch, S. Koenig, B. Kraenzlin and B. Wedemeier for technical assistance, and S. Beier for bioinformatic support with the genetic map of *T*. *aestivum*. We thank ICARDA, IPK, USDA, the Czech Crop Research Institute, and the Kyoto University Laboratory of Plant Genetics for providing seed materials. The assemblies of the 244 enriched nuclear loci (Dataset S1), the demultiplexed fasta-file of the barcoded reads for each accession used for GBS and the matrix for the filtered loci (Dataset S2) are published via e!DAL (Arend et al. 2014) at http://dx.doi.org/XXX.

