## Supplementary Information for "Genome-wide sequence information reveals recurrent hybridization among diploid wheat wild relatives"

**This PDF file includes:**

Supplementary text
Figures S1 to S10
References for SI reference citations

**Other supplementary materials for this manuscript include the following:**

**SI appendix:**
Tables S1 to S8

**SI data stored at eDAL!** (Arend et al. 2014)
Datasets S1-S2

### SI Materials and Methods

Design of capture probes and library preparation for target-enrichment.
For the development of a bait library, first, 10,105 publicly available barley full-length complementary DNAs (fl-cDNAs) from the cultivar 'Haruna Nijo' (Matsumoto et al. 2011) were retrieved from the Triticeae full-length CDS database (Mochida et al. 2009). These fl-cDNAs were assembled in a putative linear gene model employing a series of bioinformatically constructed genome zippers, that compared the fully sequenced grass genomes of *Brachypodium distachyon*, rice, and sorghum and used the extensive conservation of synteny between them (Mayer et al. 2011). ORTHOMCL (Chen et al.

2006) was used to remove loci for which orthology could not be verified between these taxa. Then one locus was selected every 0.5 cM resulting in 2,129 fl-cDNAs (approximately 300 per chromosome) that were used as a query for BLAST sequence comparison (Altschul et al. 1990) against data of *Brachypodium*, rice and sorghum, barley and wheat. If a gene was matched by the BLAST search, the genomic and fl-cDNA sequences of the corresponding gene (first best hit) were included in a multiple sequence alignment (MSA) using the MUSCLE alignment algorithm (Edgar 2004). Exon/intron boundaries were identified by the genomic DNA sequences of *Hordeum vulgare* cv. Morex and *Brachypodium distachyon* in the alignments. The loci were selected manually for bait design when the alignment met the following criteria: (i) a conserved exon-intron structure for most of the taxa considered in the alignment, (ii) a total length of exonic region to be enriched per locus larger than 1000 bp with (iii) a minimum size of single exons being 120 bp, and (iv) introns separating adjacent short exons being smaller than 400 bp. Using these criteria should enable to recover a high extent of genetic diversity by not only retrieving sequence information from exons but also from introns or parts of introns next to them. Due to higher substitution rates in introns, their sequence information would provide additional phylogenetic information to the dataset and would be of high value for understanding relationships between closely related taxa. Based on these criteria and the information on chromosomal locations, alignments of 451 loci were selected for bait design. Although the MSAs comprised also sequences from sorghum and rice, the design of capture probes was finally based only on fl-cDNAs from *H. vulgare* and *Triticum aestivum*, two distantly related Triticeae taxa, and *B. distachyon*, which was used to further consider outgroup information. Capture probes for each of the 451 loci were designed on exon sequences of all three species. Baits were designed to cover each exon at least three times. The total exonic sequence information considered in bait design amounts to 690 kb. Bait design was performed with custom PERL scripts and submitted to the web-based application eARRAY (Agilent Technologies). The loci used for bait-design are evenly disturbed over the seven chromosomes of the A, B and D-genome of *T. aestivum* (Table S1, Fig. S1).

The LE220 Focused-Ultrasonicator (Covaris) was used to shear 3 µg genomic DNA in 130 µl TE buffer for every sample into fragments having an average length of 400 bp with the following settings: instantaneous ultrasonic power (PIP) 450 W, duty factor (df) 30%, cycles per burst (cpb) 200. The treatment was applied for 100 sec. The sheared DNA was used in a sequence-capture approach (SureSelectXT Target Enrichment for Illumina Paired-End Sequencing, Agilent Technologies) targeting at 451 nuclear single-copy loci aiming for 0.01–0.02% of a Triticeae genome. All samples were barcoded and pooled (Meyer and Kircher 2010; Himmelbach et al. 2014) at equimolar ratios. Capture libraries were sequenced on the Illumina HiSeq 2000 or MiSeq. The flowcells were loaded aiming for a sequencing coverage of 40X.

<u>Target-enrichment data assembly and analyses.</u>
The assembly was performed in a two-step procedure. (i) All 451 loci considered in capture probe design were assembled in a fast and non-stringent approach to evaluate if the capture worked sufficiently and if the loci are truly single-copy in most of the taxa

selected. Therefore, the sequence assembly of the barley genome (Mayer et al. 2012) was used as a reference for the primary data analysis. Since the capture probes were designed prior to its release, the exon sequences of fl-cDNAs from *Hordeum vulgare* were first aligned to the annotated barley genome assembly using BLAT (Kent 2002) with default parameters. The physical location of an annotated exon was chosen for downstream analysis if it was the best BLAT hit and query and target sequence had an overlap of at least 60%. Exons belonging to the same gene and including introns were merged into a target locus using BEDTOOLS version 2.22.0 (Quinlan and Hall 2010). Potentially due to incomplete annotation of the barley genome assembly, some exons of fl-cDNAs did not map to annotated exons. The best BLAT hits of these exons were also kept as target regions. For each sample, the sequence reads were mapped to the barley genome assembly using the Burrows-Wheeler Alignment (BWA) Tool v. 0.7.8 (Li and Durbin 2009). The parameters k (i.e. maximum difference in seed) and n (i.e. maximum edit distance) were adjusted to allow the simultaneous mapping of reads with different lengths (MiSeq and HiSeq reads). The final mismatch rate was approximately 10%. Consensus sequences were called using SAMTOOLS version 1.1. (Li et al. 2009; Li 2011) and converted into FASTA sequences using VCFUTILS and SEQTK version 1.0 (Heng Li, https://github.com/lh3/seqtk). Consensus sequences for each of the 451 target loci were extracted. For all samples, the sequences assembled for the captured loci were subsumed in locus-wise multiple sequence alignments. The percentage of ambiguous sites was determined for each sequence in the multiple sequence alignment for each locus separately using a custom PERL script. Allelic diversity is assumed to be much lower than 1% for single- and low-copy-number loci. Thus, a high percentage of ambiguous positions for sequences of the same species are assumed to reflect the presence of paralogous gene copies as in (Jakob et al. 2014). Loci showing an average value of ambiguous sites >1% in more than five species were considered as multi-copy. Only the loci that were defined as mainly low-copy-number loci were used for further quality assessment and for phylogenetic inferences. Loci were selected if the locus had a length of at least 1000 bp, contained less than 25% of missing data and at least 15% of parsimony-informative positions, as identified with PAUP*4.0b10 (Swofford 2002).
(ii) GENEIOUS was chosen to perform a second refined assembly for quality filtered loci as it allows the reliable assembly of short insertions and deletions. Also, the assemblies can be easily visualized and necessary adjustments can be made. The sequence assembly was entirely performed in GENEIOUS v. 10.0.5 (Kearse et al. 2012), http://www.geneious.com). First, an in silico enrichment was performed by mapping all reads to the sequence of the whole-chloroplast genome sequence of *Hordeum vulgare* (EF115541). The unused reads were stored and reads were deduplicated using Dedupe Duplicate Read Remover 36.32 (as part of BBMap - Bushnell B. - sourceforge.net/projects/bbmap/) with default settings in GENEIOUS. Then, regions with more than a 5% chance of an error per base were trimmed (error probability limit). The genomic sequence of loci defined as mainly single-copy in previous steps was extracted from the genomic sequence of *Hordeum vulgare* cv. Morex to be used as a reference for mapping. Only paired reads that mapped nearby were mapped to the reference sequences of the enriched locus with 10% as a maximum of mismatches per read. Further, the GENEIOUS assembly algorithm was allowed to find deletions of up to 200 bp and indels

with an insert size of up to 350 bp. The mapping quality was set so that a read is mapped with a probability of 90% the mapping is correct.
A consensus sequence was called using the 50% majority rule, which calls bases matching at least 50% of the sequences. An "N" was called for regions of the assembly having a coverage lower than 3. The consensus sequence was trimmed to the reference sequence. Sequences aligned per locus using MAFFT version 7.3.05 with "N" treated as a wildcard on the Galaxy instance of the IPK (http://galaxy.ipk-gatersleben.de/; Giardine et al. 2005). In a second step "N"s were replaced by gaps and the alignments were realigned with MAFFT in GENEIOUS for refinement. Regions with 75% of missing data were masked before using the alignments for phylogenetic inferences.

Identification of hybrid taxa.

Four-Taxon *D* statistic tests were performed using the routine DTRIOS of DSUITE (Malinsky 2019; https://github.com/millanek/Dsuite). We first tested if *Taeniatherum caput-medusae* was involved in any introgressions by using *Dasypyrum villosum* as outgroup. We also provided the TETRAD topology (Fig. 1B; S8) inferred from the GBS data to specify species relationships. *D* statistics significance was assessed using jackknife (Green et al. 2010) on blocks of 100 SNPs. The function *p.adjust* in R 3.5.3 (R Core Team 2019) was used to apply a Benjamini-Yekutieli correction (Benjamini and Yekutieli 2001). The analysis generated 320 tests summarized and visualized with the Ruby script "plot_d.rb" available from M. Matschiner (https://github.com/mmatschiner).

### Supplementary Figures

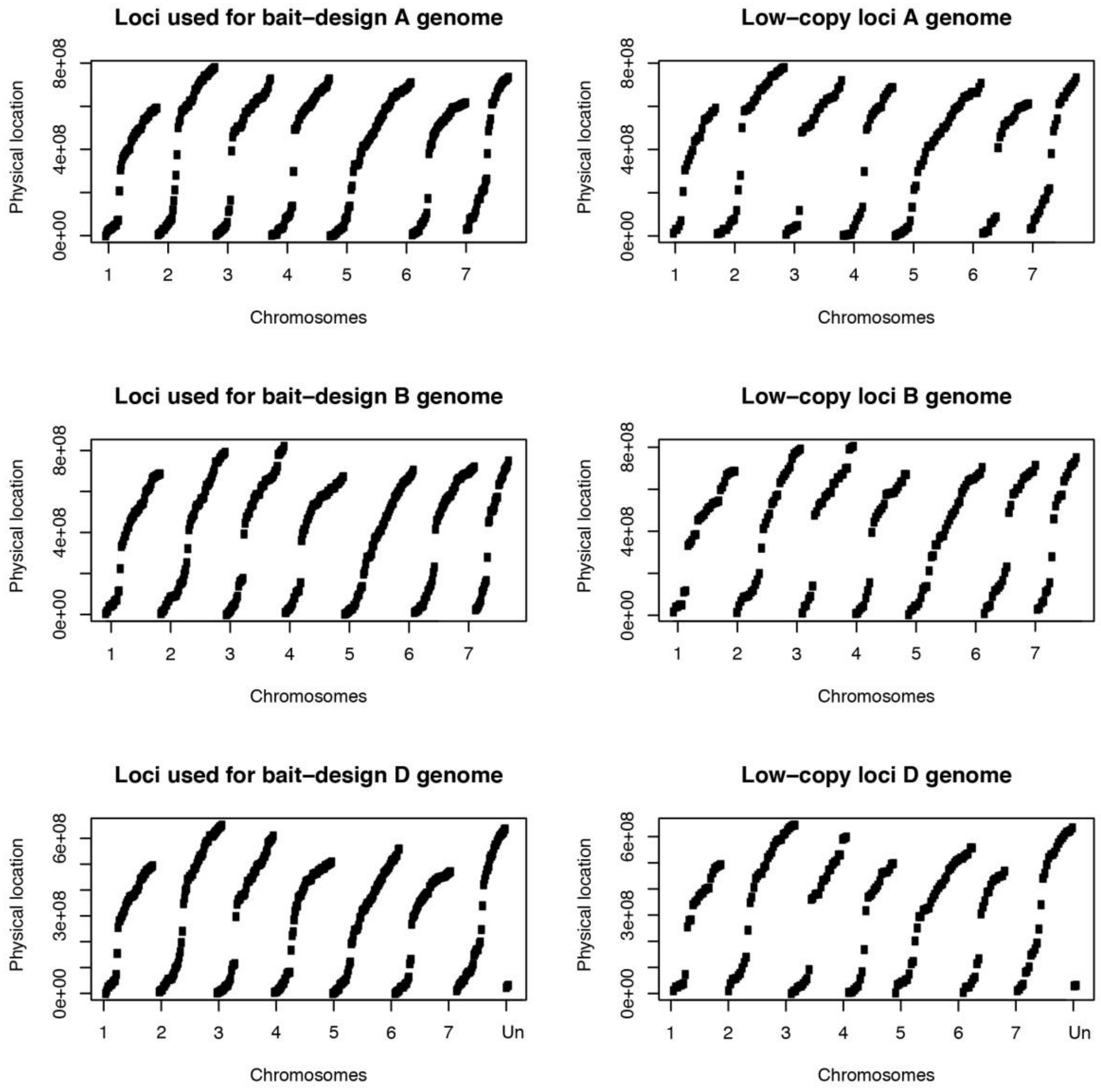


**Figure S1. Representation of the physical locations of loci considered in bait design (left side) and low-copy loci used for analyses (right side) in the** A-**,** B- **and** D**-genome of *T. aestivum*.** The barley fl-cDNA used initially for the selection of target loci were mapped against the seven chromosomes of each subgenome of *T. aestivum* (International Wheat Genome Sequencing Consortium 2018) using GMAP version 2018-03-25 (Wu and Watanabe 2005). Three loci were localized on unmapped contigs (Un).

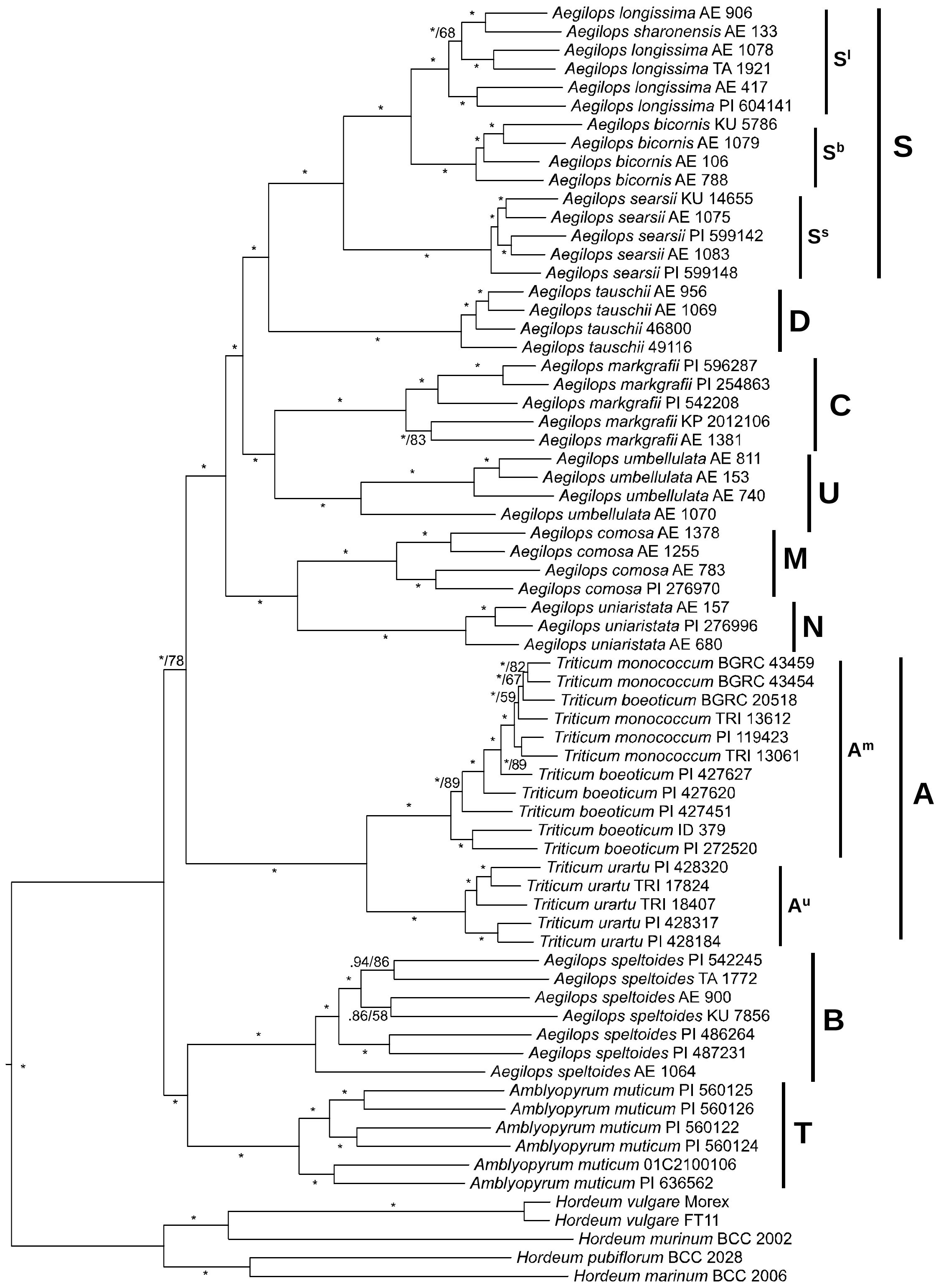


**Figure S2. Phylogenetic tree derived from Bayesian phylogenetic inference of the concatenated datamatrix of 244 loci.** The same relationships were obtained by a maximum likelihood analysis. Numbers along branches provide Bayesian posterior probabilities (pp)/ML bootstrap support (bs) values. Asterisks indicate a pp of 1/bs ≥90%. Genome names are given to the right.

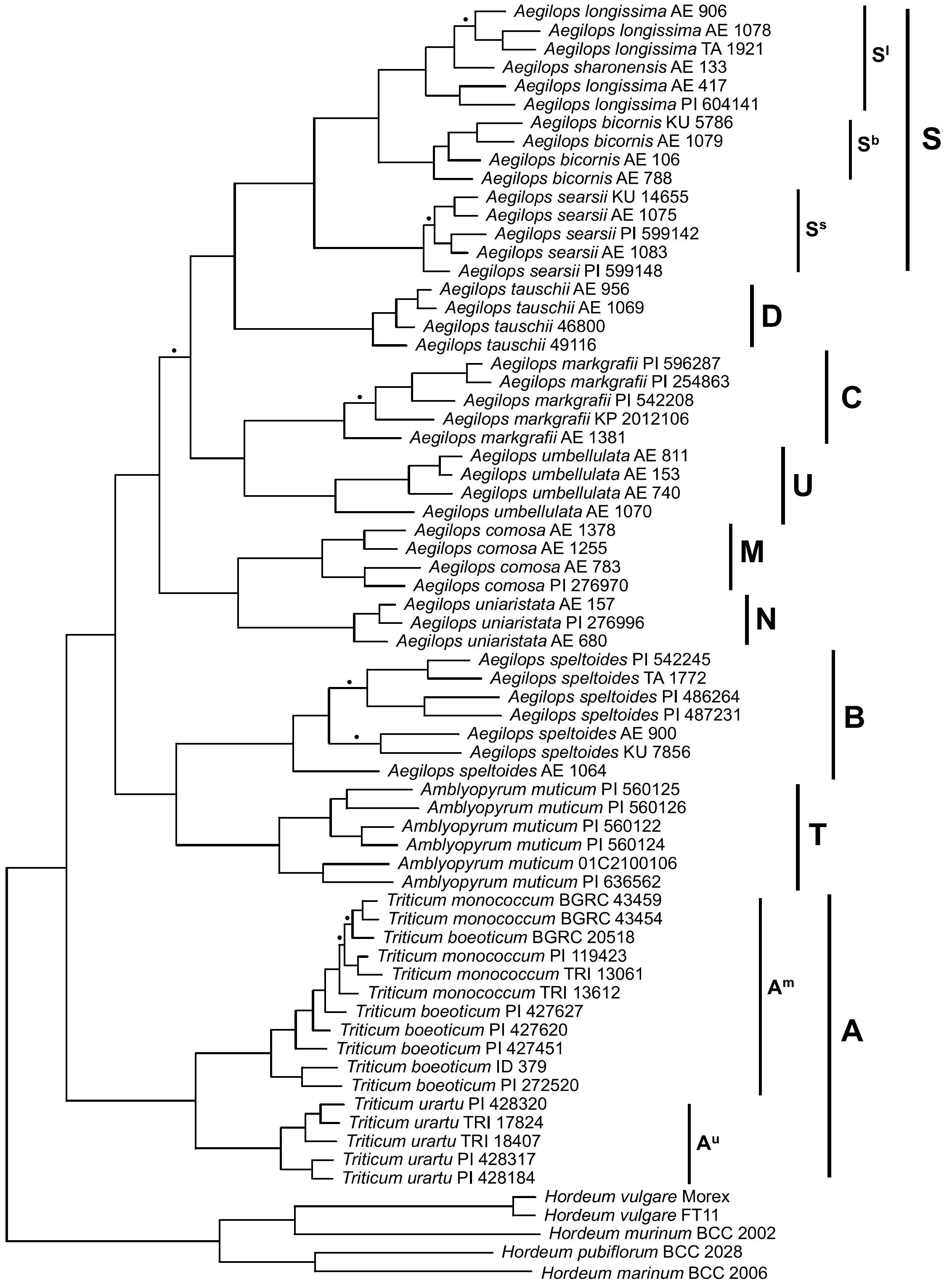


**Figure S3. Single maximum parsimony tree derived from the concatenated datamatrix of 244 loci.** Bullet points indicate branches where bootstrap support values were below 70%. For the unmarked branches support values are ≥96%, mostly 100%. Genome names are provided to the right.

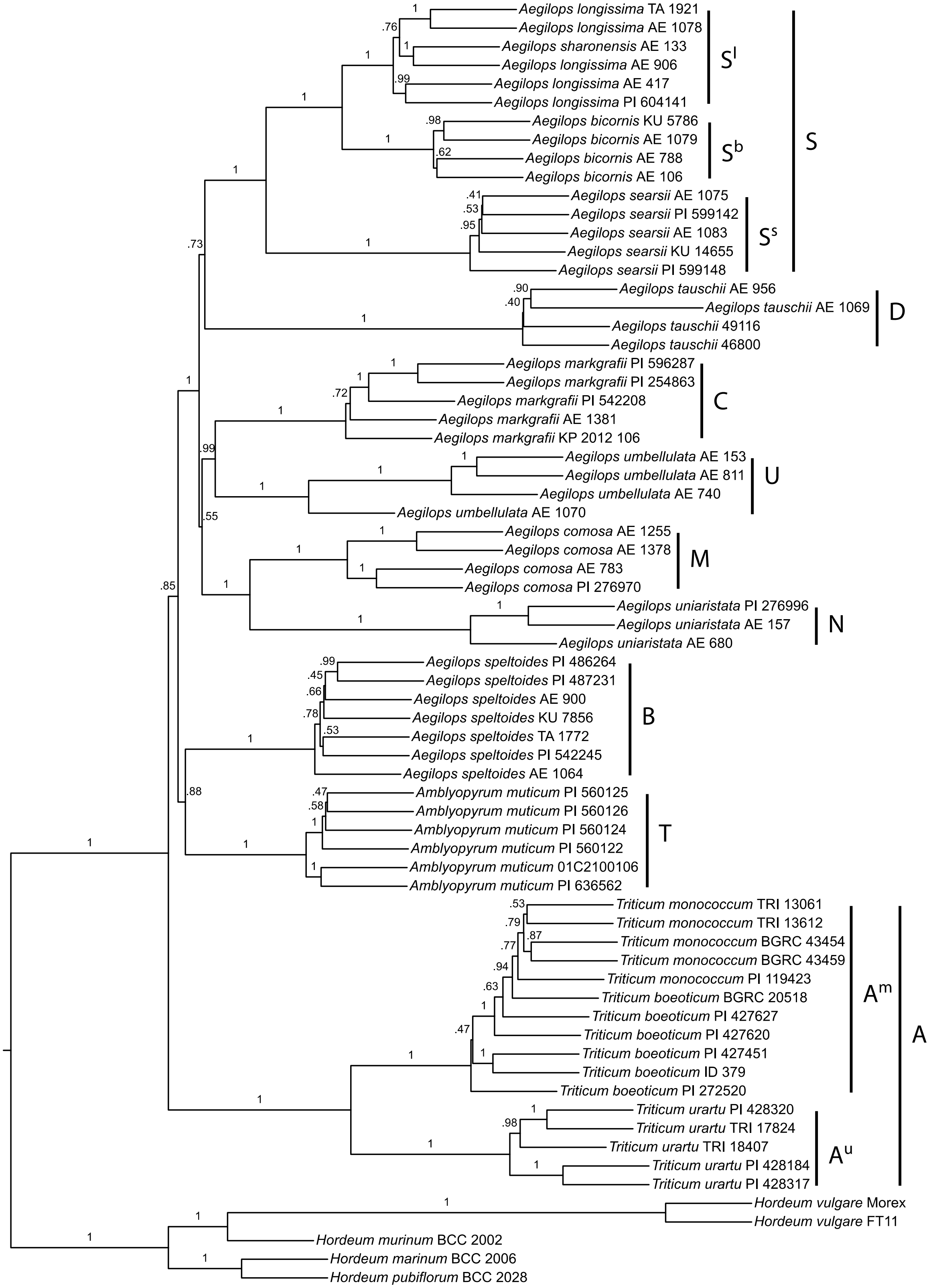


**Figure S4. Multispecies coalescent tree calculated with ASTRAL from separate ML trees of 244 loci.** Numbers along branches provide local posterior probability values. Genome names are given to the right.

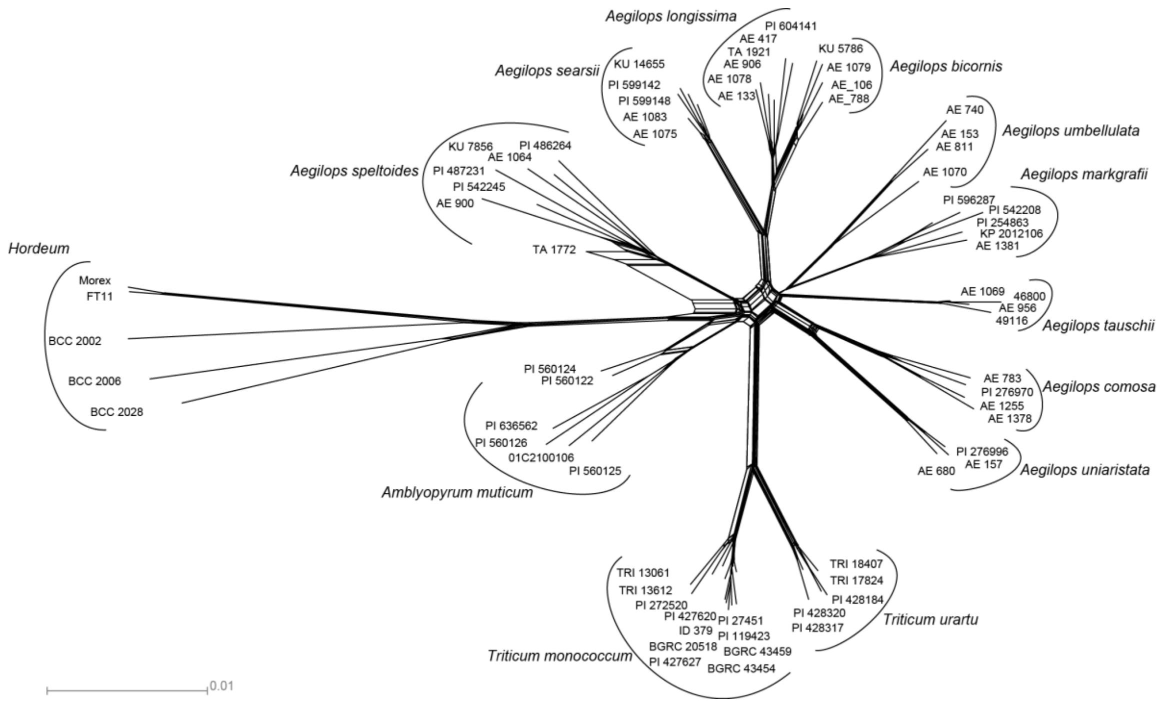


**Figure S5. Phylogenetic network inferred via the NEIGHBORNET method in SPLITSTREE4 from the concatenated matrix of 244 target enriched loci.**

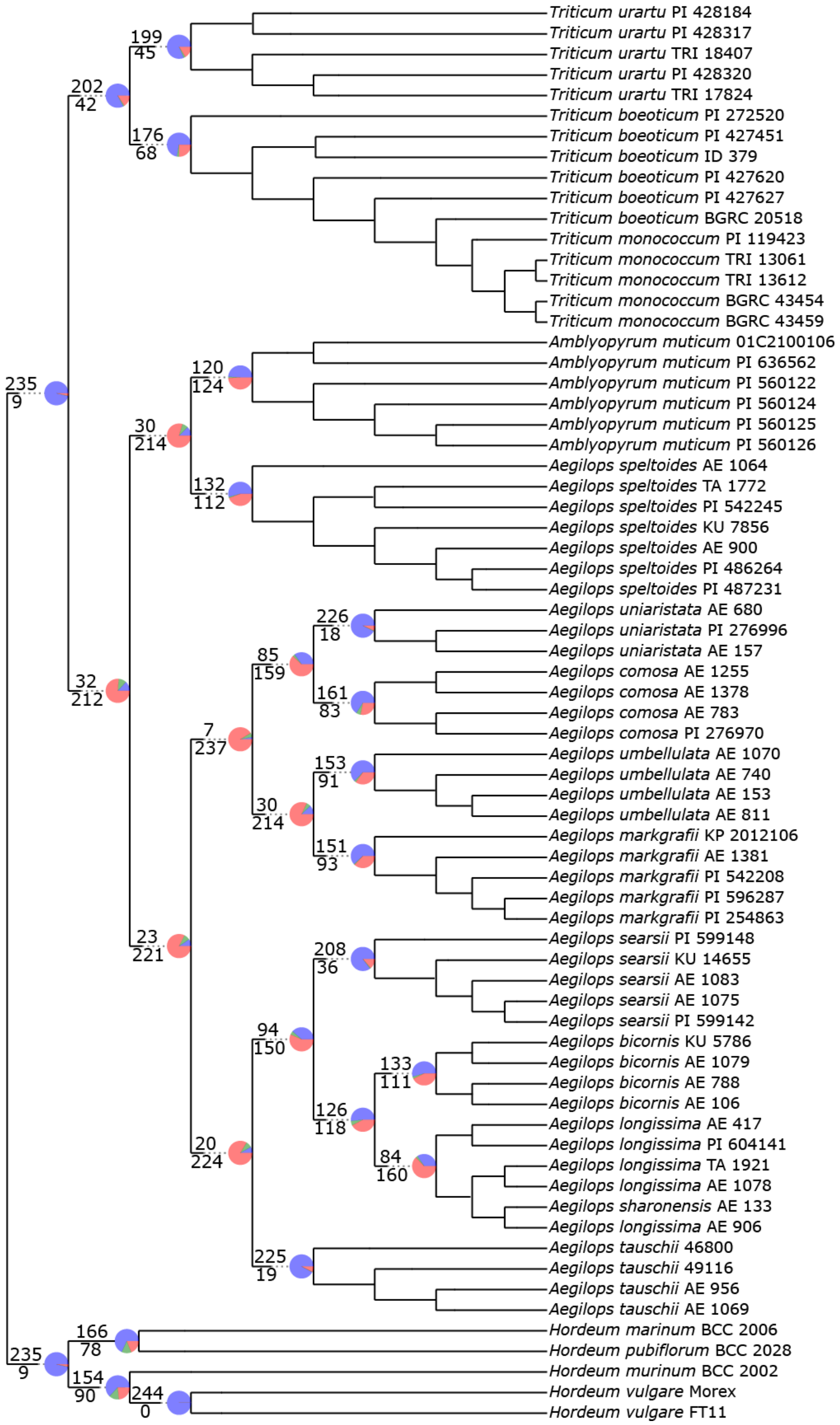


**Figure S6 Visualization of the degree of gene tree/species tree conflict**. Conflict among the individual ML gene trees in comparison to the ASTRAL species tree was investigated using PHYPARTS (Smith et al. 2015). Gene tree/species tree conflicts are reported only for single and multispecies clades with the number on top representing the number of gene trees concordant with the species tree, while the number below indicates the number of topologies in conflict with that clade in the species tree. Pie charts show the proportion of gene trees supporting a clade (blue), the main alternative for that clade (green), and the proportion supporting the remaining alternatives (red).

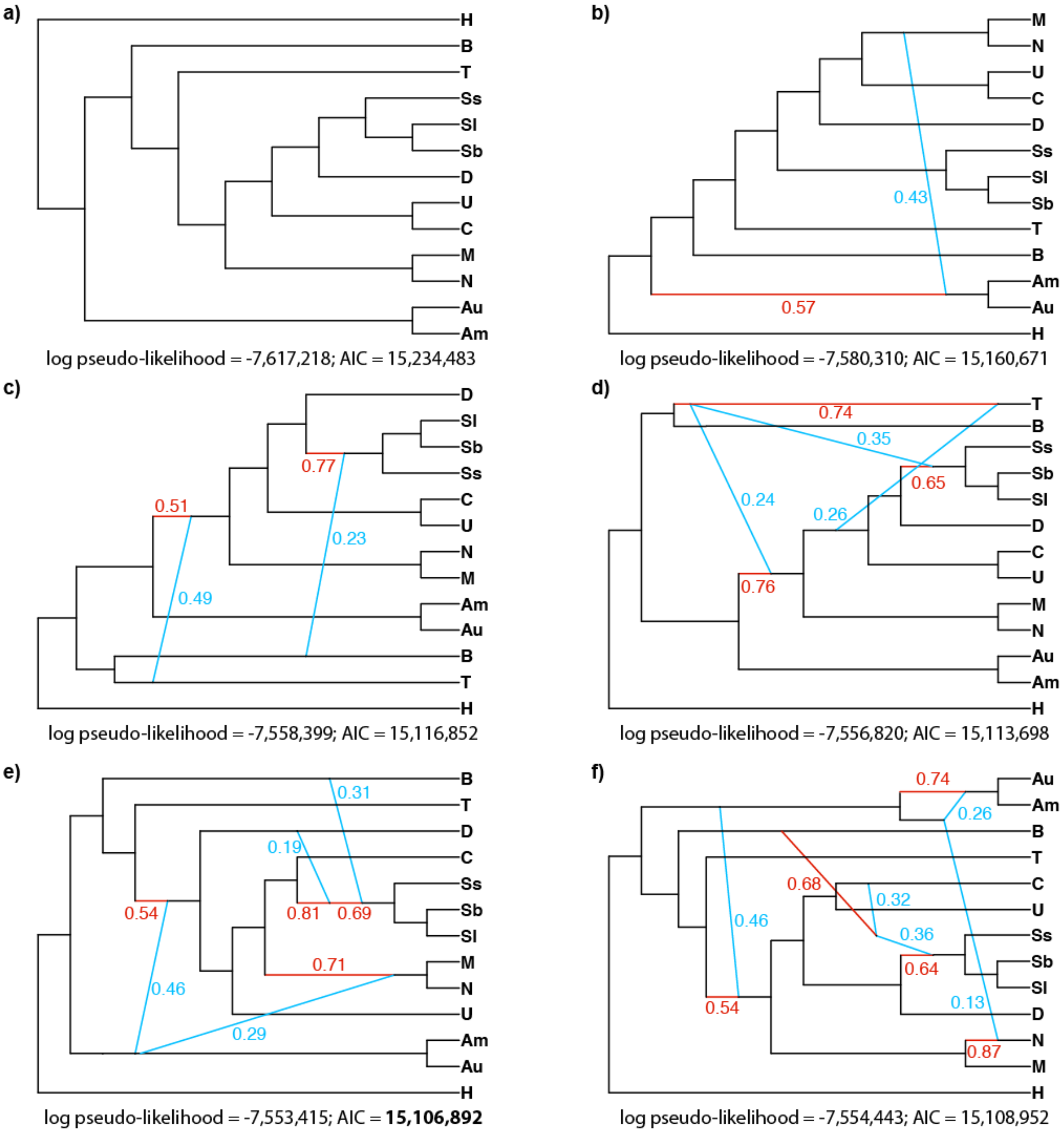


**Figure S7 Phylogenetic networks inferred under the multispecies network coalescent (MSNC) from the 244 gene tree topologies with maximum pseudo-likelihood.** Networks a-f were inferred with the routine InferNetwork_MPL of PhyloNet setting a maximum of zero to five reticulations, respectively. Major genome contribution to hybrid lineages are indicated in red and introgressions are indicated in blue. Numbers represent estimated inheritance probabilities. The networks were visualized with PhyloNetworks (Solís-Lemus et al. 2017).

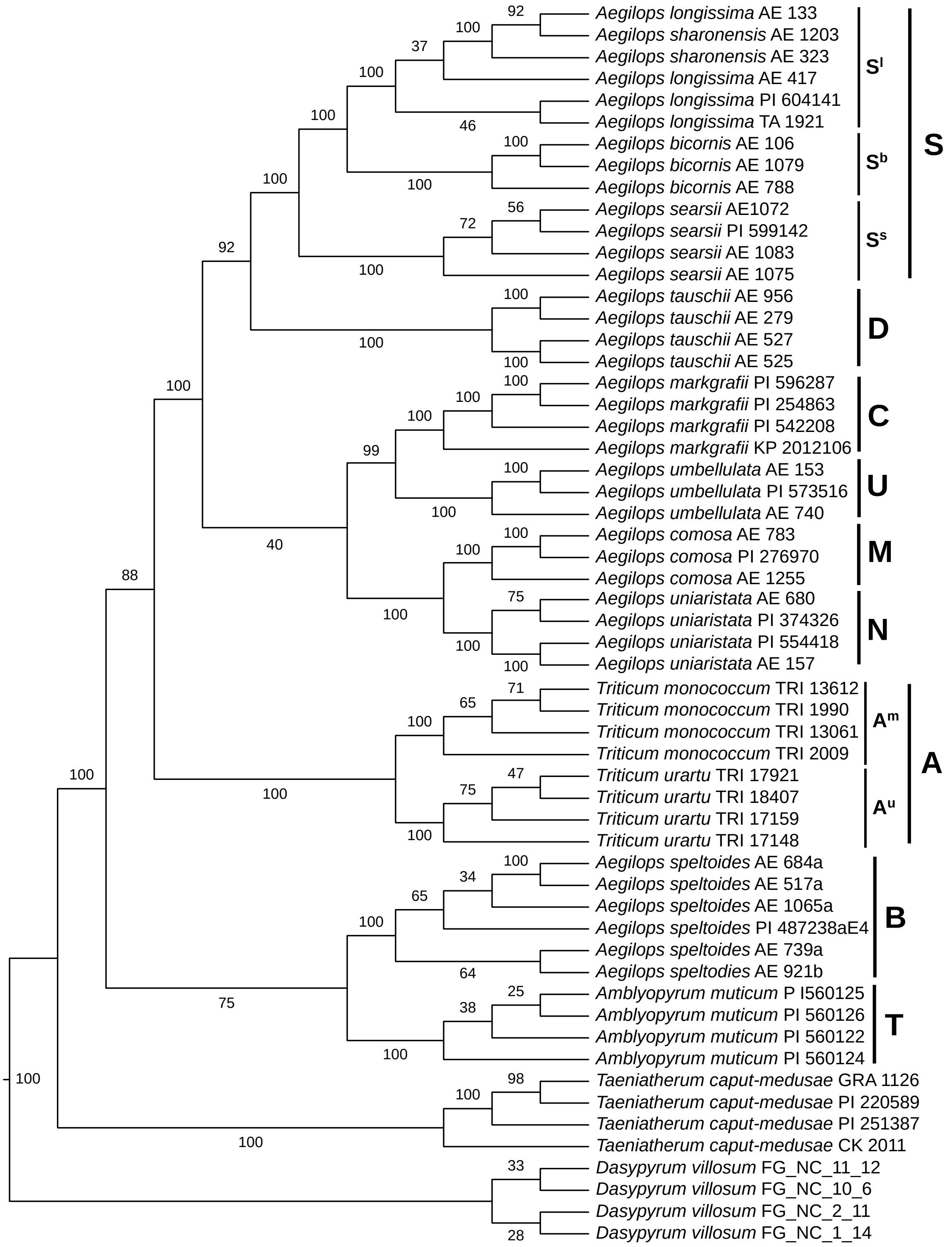


**Figure S8. Consensus cladogram derived from a TETRAD analysis of the GBS data.** Numbers along branches are TETRAD bootstrap values (%). As *Hordeum* is taxonomically too remote from the wheat group species to be used as outgroup in a GBS-based analysis, we included *Dasypyrum* and *Taeniatherum* instead. Genome designations for the species are given to the right.

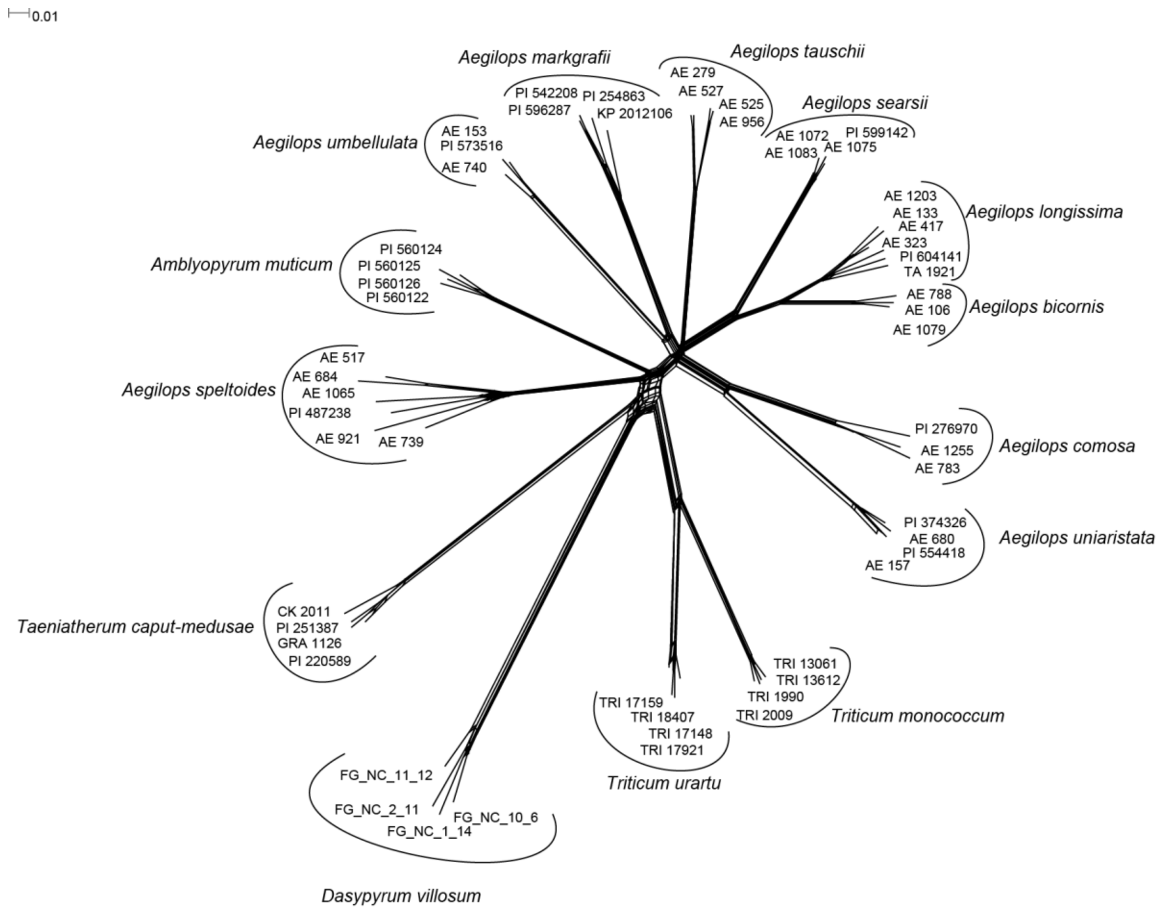


**Figure S9 Phylogenetic network inferred via the NEIGHBORNET method in SPLITSTREE4 from 807,909 SNPs using the full GBS matrix.**

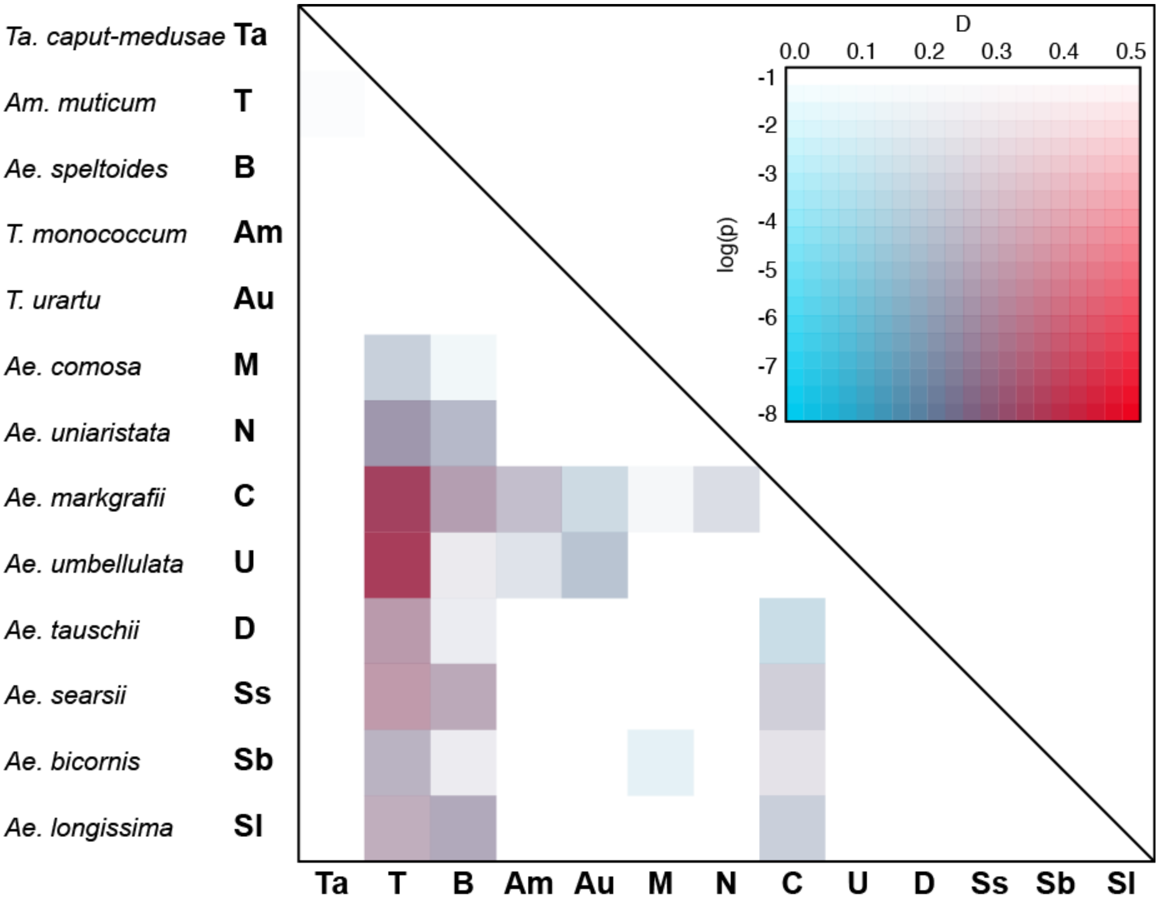


**Figure S10. Heatmap summarizing Four-taxon *D* statistic tests using *Dasypyrum villosum* as outgroup**. The plot is based on 286 tests. It shows the *D* statistic results and their significance for each pair of species. Red and blue indicate high and low *D* statistic values, respectively. The intensity of the color corresponds to the *p* value (in log-scale) assessed using the block jackknife procedure and corrected with Benjamini-Yekutieli for multiple testing.

**Additional data Table S1 (separate file)**

**Overview of loci selected for the sequence capture.** The locus name refers to the barley fl-cDNA initially used for loci selection (Mayer et al. 2012). Physical locations of these loci in bread wheat were obtained using the most recent genetic map of bread wheat(International Wheat Genome Sequencing Consortium 2018).

**Additional data Table S2 (separate file)**

Read statistics of the target-enrichment experiment.

**Additional data Table S3 (separate file)**

Table 3a Percentages of ambiguous positions per sequenced accession and locus as estimated by a custom PERL script.
Table 3b The average percentages of ambiguous positions per locus were compared for every genus. Loci showing an average value of ambiguous sites >1% in more than five diploid species were considered as mainly multi-copy loci.

**Additional data Table S4 (separate file)**

Table S4a. Overview of the number of reads mapping and the medium coverage for each of the 244 loci and each accession. For each locus the length of the genomic reference sequence from barley is given.
Table S4b. Information on the multiple sequence alignments used for phylogenetic inference.

**Additional data Table S5 (separate file)**

**Overview of the material considered in this study.** For all materials, the seed bank identifier (if existing) and the species name as used in this study, as well as their species synonyms used in the donor seed banks (Material source), are provided. The genome symbol and the country of origin, where the material was originally collected, are given. The information if a herbarium voucher could be deposited in the herbarium of IPK Gatersleben (GAT) is given. The last two columns show the kind of data that was generated for the individual accessions.

**Additional data Table S6 (separate file)**

Read and assembly statistics for the GBS data.

### Additional data Table S7 (separate file)

**Four-taxon *D* statistics.** Three ingroup and one outgroup taxa were tested using a fixed phylogeny (((P1, P2) P3), O), with P3 being the focal taxon. All tests were done with *Taeniatherum caput-medusae* as outgroup (O). The ASTRAL topology (Fig. 1A) was used to specify species relationships. *p-values* were assessed using the block jackknife procedure and adjusted using a Benjamini-Yekutieli correction (adj_p-value). A cutoff of 0.05 was then used for significance.

**Additional data Table S8 (separate file)**

**$D_{FOIL}$ test for introgression in wheat wild relatives from GBS data.** For each test, the combination of species, identified by their genome denomination (cf. Table S2), is indicated as P1-P4. All tests were done with *Taeniatherum caput-medusae* as outgroup. For each test, the total number of sites is indicated (total) as well the number of patterns analyzed (dtotal). The score of the test (Dxx_stat), the Chi-squared value (Dxx_chisq) and associated P-value (Dxx_Pvalue) are reported (see Pease and Hahn 2015 and $D_{FOIL}$ manual for details). Warnings raised by the program are reported: "discordant patterns are more frequent than concordant ones" (a), "divergences are not ordered correctly" (b) and "terminal branch pairs are not proportionate" (c). * indicate tests performed with dfoilalt mode to correct for "an accelerated substitution rate on a specific branch relative to its sister taxon". Single arrows (→) indicate the direction of introgression while double arrows (<=>) indicate no direction inferred. Red background cells indicate relationships that raised a "b" warning and that have been ignored as potentially misleading. *p-values* were adjusted using a Benjamini-Yekutieli correction (adj_Dxx_Pvalue). A cutoff of 0.01 was then used for significance.
